# Strand-resolved mutagenicity of DNA damage and repair

**DOI:** 10.1101/2022.06.10.495644

**Authors:** Craig J. Anderson, Lana Talmane, Juliet Luft, Michael D. Nicholson, John Connelly, Oriol Pich, Susan Campbell, Vasavi Sundaram, Frances Connor, Paul A. Ginno, Liver Cancer Evolution Consortium, Núria López-Bigas, Paul Flicek, Colin A. Semple, Duncan T. Odom, Sarah J. Aitken, Martin S. Taylor

**Affiliations:** Medical Research Council Human Genetics Unit, Institute of Genetics and Cancer, University of Edinburgh, Edinburgh, UK; Institute of Genetics and Cancer, University of Edinburgh, Edinburgh, UK; Medical Research Council Toxicology Unit, University of Cambridge, Cambridge, UK; Edinburgh Pathology, Institute of Genetics and Cancer, University of Edinburgh, Edinburgh, UK; Laboratory Medicine, NHS Lothian, Edinburgh, UK; Institute for Research in Biomedicine (IRB Barcelona), The Barcelona Institute of Science and Technology, Barcelona, Spain; European Molecular Biology Laboratory, European Bioinformatics Institute, Hinxton, UK; Cancer Research UK Cambridge Institute, University of Cambridge, Cambridge, UK; German Cancer Research Center (DKFZ), Heidelberg, Germany; Universitat Pompeu Fabra (UPF), Barcelona, Spain; Institució Catalana de Recerca i Estudis Avançats (ICREA), Barcelona, Spain; Department of Genetics, University of Cambridge, Cambridge, UK; Department of Pathology, University of Cambridge, Cambridge, UK; Department of Histopathology, Cambridge University Hospitals NHS Foundation Trust, Cambridge, UK

## Abstract

DNA base damage is a major source of oncogenic mutations^1^. Such damage can produce strand-phased mutation patterns and multiallelic variation through the process of lesion segregation^2^. Here, we exploited these properties to reveal how strand-asymmetric processes, such as replication and transcription, shape DNA damage and repair. Despite distinct mechanisms of leading and lagging strand replication^3,4^, we observe identical fidelity and damage tolerance for both strands. For small DNA adducts, our results support a model in which the same translesion polymerase is recruited on-the-fly to both replication strands, starkly contrasting the strand asymmetric tolerance of bulky adducts^5^. We find that DNA damage tolerance is also common during transcription, where RNA-polymerases frequently bypass lesions without triggering repair. At multiple genomic scales, we show the pattern of DNA damage induced mutations is largely shaped by the influence of DNA accessibility on repair efficiency, rather than gradients of DNA damage. Finally, we reveal specific genomic conditions that can corrupt the fidelity of nucleotide excision repair and actively drive oncogenic mutagenesis. These results provide insight into how strand-asymmetric mechanisms underlie the formation, tolerance, and repair of DNA damage, thereby shaping cancer genome evolution.

## Main text

There is an elegant symmetry to the structure and replication of DNA, where the two strands separate and each serves as a template for the synthesis of new daughter strands. Despite this holistic symmetry, many activities of DNA are strand asymmetric: (i) during replication different enzymes mainly synthesise the leading and lagging strands^3,4,6,7^, (ii) RNA transcription uses only one strand of the DNA as a template^8^, (iii) one side of the DNA double helix is more associated with transcription factors^9^, and (iv) alternating strands of DNA face towards or away from the nucleosome core^10,11^. These processes can each impart strand asymmetric mutational patterns that reflect the cumulative DNA transactions of the cells in which the mutations accrued^1,8–10,12^.

Cancer genomes are the result of diverse mutational processes^1,13^, often accumulated over decades, making it challenging to identify and subsequently interpret their relative roles in generating spatial and temporal mutational asymmetries. The relative contribution of DNA damage, surveillance, and repair processes to observed patterns of mutational asymmetry remains poorly understood, though mapping of DNA damage^14–17^ and repair intermediates^18,19^ have provided key insights.

We recently discovered DNA lesion segregation, a pervasive mechanism of DNA damage induced mutagenesis, which is ubiquitous for all tested mutagens in human cells and a feature of human cancers^2^. Lesion segregation was first identified in murine liver tumour genomes initiated by a single burst of chemical mutagenesis^2^. The resulting mutations had pronounced strand asymmetry, indicating that most mutagenic DNA lesions persist through at least one round of DNA replication before segregating into daughter cells (**Fig. 1a**). Since almost all (>96%) of these mutations arise from DNA lesions induced in a discrete burst of damage within a single cell-cycle, in each tumour we could identify the lesion-containing strand across half of the autosomal genome and the entire X chromosome (**Fig. 1b**). Here, we newly exploit these strand-resolved lesions as a powerful tool to quantify how mitotic replication, transcription, and DNA-protein binding mechanistically shape DNA damage, genome repair, and mutagenesis.

**Fig.1.**
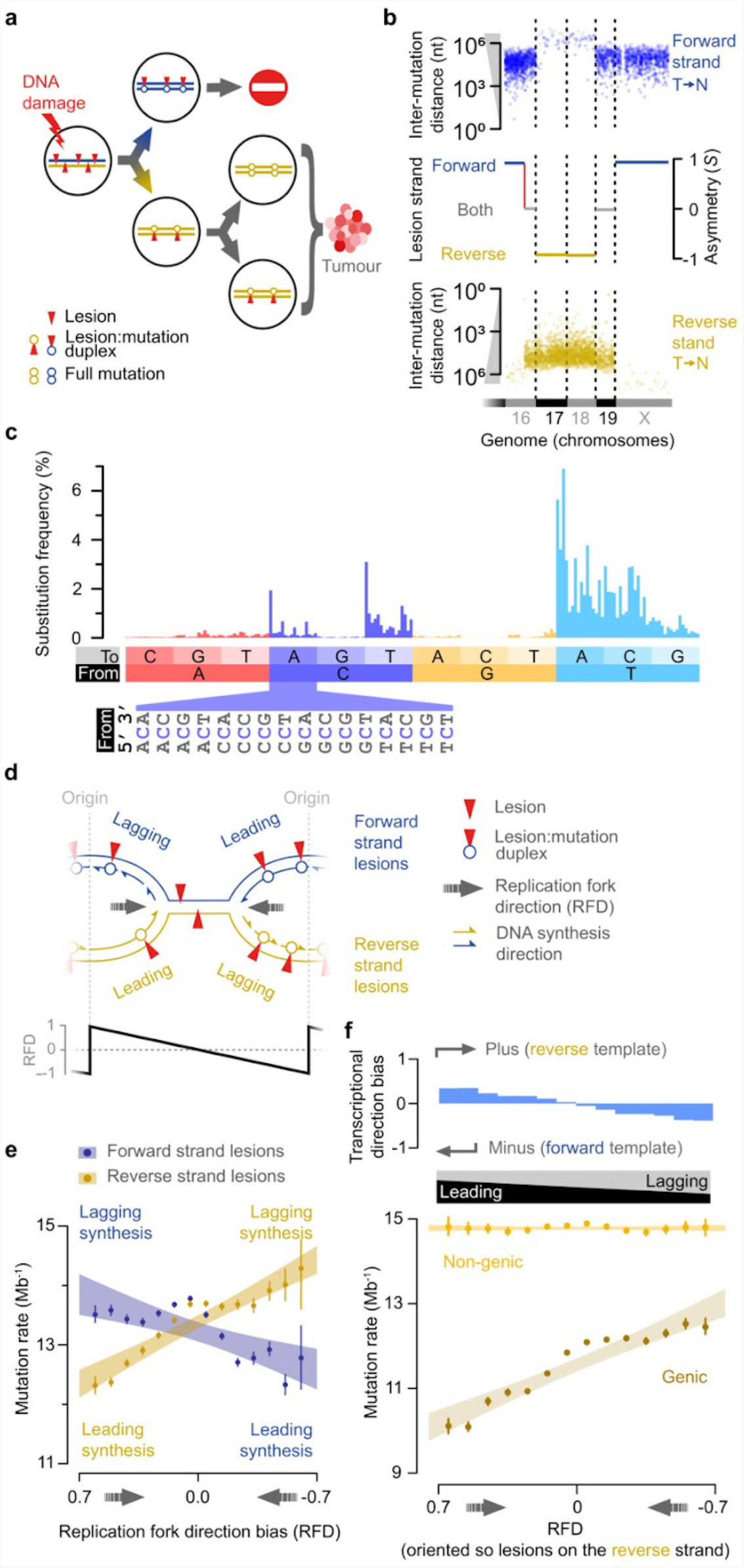
Replication strand mutational asymmetry can be explained by transcription coupled repair. **a**, DNA lesion segregation on one haploid chromosome ^2^. Mutagen exposure induces lesions (red triangles) on both DNA strands (forward blue, reverse gold). Unrepaired lesions that persist until replication serve as a reduced fidelity template. The two sister chromatids segregate into distinct daughter cells at mitosis, so new mutations are not shared between daughter cells of the first division. Since only one daughter lineage transforms and clonally expands, all damage-induced mutations in the tumour arise from lesions on only one of the originally damaged DNA strands. **b**, Mutational asymmetry in an exemplar tumour (94315_N8) segmented to resolve the lesion strand. Partial genome span (x-axis) showing individual mutations from T on the forward strand (blue) or T on the reverse strand (gold), and distance to neighbouring T mutation (y-axis). Segmentation track shows blocks of consensus asymmetry; red line shows asymmetry switches due to sister chromatid exchange. **c**, 192 category unfolded mutation spectra of all tumours (n=237), representing the relative frequency of strand-specific single-base substitutions. **d**, During the first DNA replication after DNA damage, template lesions (red triangles) are encountered by both the extending leading and lagging strands. **e**, Higher mutation rate for lagging strand than leading strand synthesis over a DNA lesion containing template. Replication fork direction (RFD, x-axis) indicates the bias towards leading or lagging strand replication. Mutation rates (y-axis) calculated in RFD bins (0.1 units) separately for forward and reverse strand lesions; point estimates (circles) with 95% bootstrap C.I. (whiskers); shaded area shows 95% C.I. of linear regression through the point estimates. **f**, Replication strand differences in mutation rate can be explained by transcription coupled repair. Lower panel as for **e**, with the genome partitioned into genic and non-genic fractions, and all genomic segments oriented so lesions are on the reverse strand. Transcriptional direction bias calculated as relative difference for highly expressed genes in P15 mouse liver.

To decipher strand specific DNA transactions, we analysed over seven million high confidence, lesion strand-resolved mutations called from 237 DEN-induced tumour genomes^2^. The lesion strand-resolved mutation signature (**Fig. 1c**), shows that most (>75%) of the mutations are from T nucleotides on the lesion containing strand, consistent with biochemical evidence of frequent mutagenic DEN adducts on T^20^ and prior analyses of DEN induced tumours^2,21^.

### Leading and lagging strand mutagenesis is symmetrical

These well-powered and experimentally controlled *in vivo* data provide a unique opportunity to evaluate whether DNA damage on the replication leading strand results in the same rate and spectrum of mutations as on the lagging strand template. There are several reasons why they might differ. First, leading and lagging strand replication uses distinct replicative enzymes^3,4,6,7^, which may differ in how they handle unrepaired DEN-induced damage, potentially resulting in different misincorporation profiles. Second, it is unknown whether the leading and lagging strand polymerases recruit different translesion polymerases, which could generate distinct error profiles. Third, substantially longer replication gaps are expected on the leading strand, if there is polymerase stalling^22^. Consequently, leading and lagging strands are thought to differ in their lesion bypass^5^ and post-replicative gap filling^23,24^.

Based on measures of replication fork directionality^25^ and patterns of mutation asymmetry, we inferred whether the lesion-containing strand preferentially templated the leading or lagging replication strand (**Fig. 1d**). This was separately resolved for each genomic locus on a per tumour basis. Our initial analysis demonstrated a significantly higher mutation rate for lagging strand synthesis over a lesion containing template (Pearson’s correlation coefficient cor=-0.86, p=3.2×10^−9^; **Fig. 1e**). However, gene orientation -and thus the directionality of transcription - also corresponds to replication direction^26,27^ and DEN lesions are subject to transcription coupled repair (TCR)^2^. After accounting for TCR there is no residual effect of replication strand on mutation rate (Pearson’s correlation coefficient, non-genic regions cor=0, p=0.99; **Fig. 1f**).

Given that the leading and lagging strands are synthesised by distinct cellular machinery^4,7,27– 29^, the observed consistency in the rate and spectrum of mutations generated by DEN lesions is unexpected. These results point to a shared mechanism of lesion bypass, strongly suggesting the same translesion polymerases are recruited by both the leading and lagging strands replication machineries.

### Mutation clusters demonstrate the replicative symmetry of translesion synthesis

It has been proposed that when translesion polymerases replicate across damaged bases, they can generate proximal tracts of low fidelity synthesis^30–32^. In bacteria and yeast this produces clusters of mutations^33,34^ and such collateral mutagenesis was recently reported in vertebrates^35^. Consistent with these models, we found that mutations within 10 nucleotides of each other are significantly elevated over background expectation (two tailed Fisher’s test, odds ratio 11.9, p <2.2×10^−16^). This enrichment is most pronounced for 1 to 2 nucleotide spacing, decreases abruptly after one DNA helical turn (ca. 10 nucleotides), and decays to background within 100 nucleotides (**Fig. 2a**; **Extended Data Fig. 1d-g**). These short clusters are overwhelmingly isolated pairs of proximal mutations (98% pairs, 2% trios) phased on the same chromosome (**Fig. 2b**).

**Fig.2.**
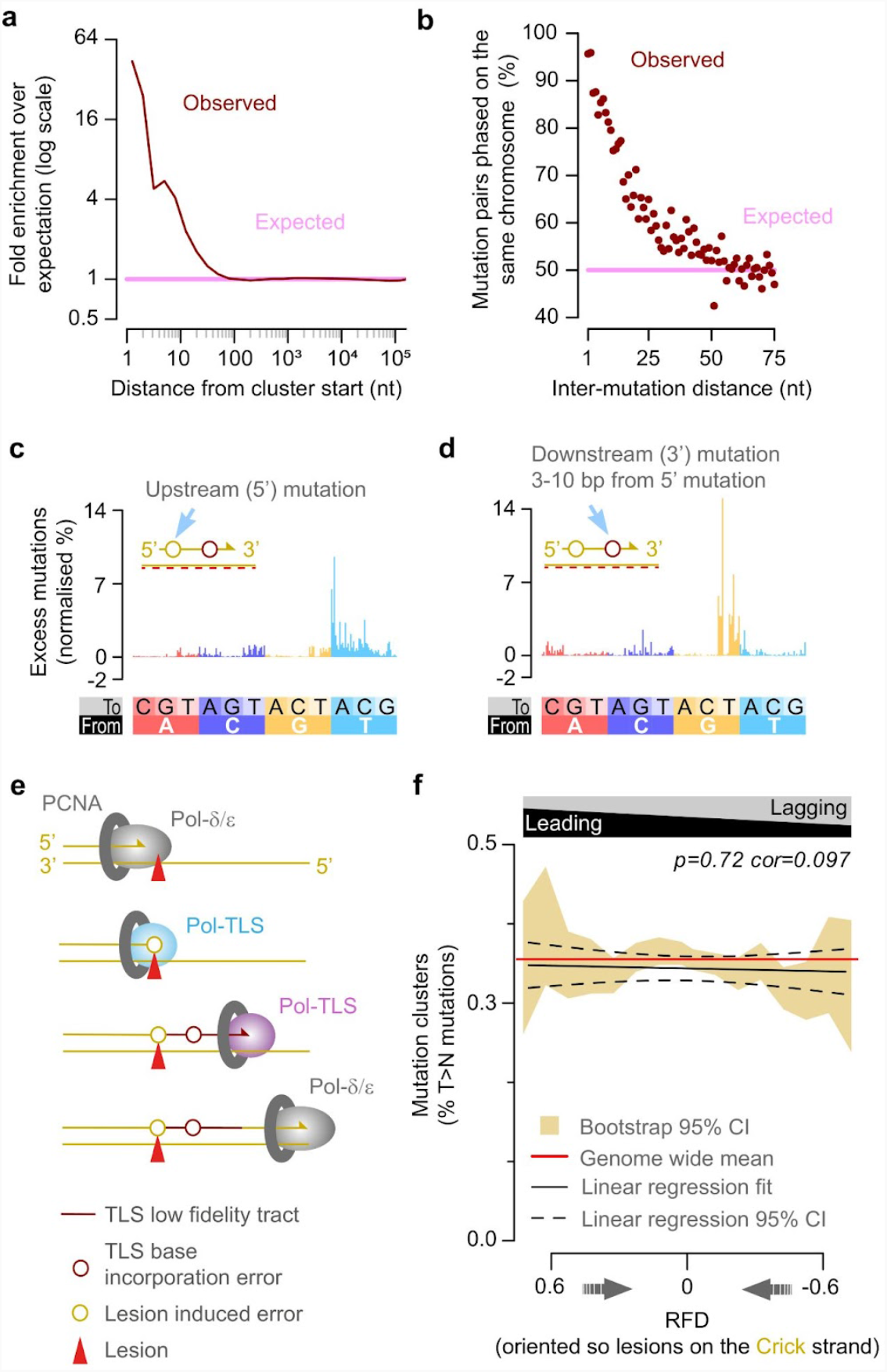
Translesion synthesis (TLS) drives collateral mutagenesis on both the leading and lagging strands. **a**, Closely spaced mutations (brown) occur more frequently than expected (pink). **b**, Clustered mutation pairs co-occur in the same sequencing read, confirming they are on the same DNA duplex. **c**, Residual mutation signature (after subtracting expected mutations) for cluster upstream mutations. Cluster orientation by the lesion containing strand (red-dash line), absolute area of histogram sums to 100. **d**, Residual signature of downstream (+3 to +10 nt) cluster mutations, plotted as per **c. e**, Schematic illustrating a replicative polymerase stalling at a template lesion (red triangle), recruiting a TLS polymerase, and inducing an upstream mutation (yellow circle). It then either extends or hands over to a second reduced fidelity polymerase, resulting in a proximal downstream mutation (collateral mutagenesis; brown circle). **f**, Mutation clusters occur consistently at 0.35% of T>N mutations irrespective of replication strand bias, indicating a similar rate of TLS for both the leading and lagging strand; there is no significant correlation between replication strand and the rate of mutation clusters.

We oriented the clusters by their lesion containing strand, and designated the first mutation site to be replicated over on the lesion containing template as the upstream (5’) mutation and subsequent mutations were designated downstream (3’). Upstream mutations showed a mutation spectrum closely resembling the tumours as a whole (**Fig. 2c**; **Extended Data Fig. 1a,b,i**), indicating that it represents a typical lesion-templated substitution.

In contrast, downstream mutations have distinct mutation spectra (**Extended Data Fig. 1c**). Those located >2 nucleotides downstream show a strong preference for G→T substitutions (**Fig. 2d**; **Extended Data Fig. 1h,l-n**). Since mutations are called relative to the lesion containing template strand, this indicates the preferential misincorporation of A nucleotides opposite a template G nucleotide, thus revealing the intrinsic error profile of an extending translesion polymerase. Mutation pairs with closer spacing (≤2 nucleotides) exhibit somewhat divergent mutation signatures (**Extended Data Fig. 1h,j,k**), likely reflecting both composition constraints and other processes such as the transition between alternate translesion polymerases (**Fig. 2e**).

Three lines of evidence supported a model in which the same translesion polymerases are recruited with equal efficiency and processivity to both the leading and lagging strands: (i) The leading and lagging strands had essentially identical relative-rates of mutation clusters (**Fig. 2f**), (ii) the mutation spectra of the downstream mutations were the same (**Extended Data Fig. 1o**), and (iii) the length distribution of clusters matched between leading strand and lagging strand biassed regions (no significant difference in size distribution, Kolmogorov-Smirnov test (p=0.98) despite >98% power to detect a difference in the distribution of cluster lengths in which ≥4.8% of ≤10 nt clusters become >10 nt clusters, **Extended Data Fig. 1p**,**q**).

Having established the replicative symmetry of damage induced mutagenesis and determined the relative contributions of replication and transcription on mutation rate, we next probed the strand-specific effects of transcription on DNA repair and mutagenesis.

### Enhanced repair of both strands in transcriptionally active open chromatin

We previously demonstrated that repair of DEN induced lesions on the transcription template strand is expression dependent^2^ (**Fig. 3a**), and have now shown that most replication strand asymmetry in both mutation rate and spectrum can be attributed to transcription, rather than replication (**Fig. 1f**). Nascent transcription estimates, as expected^36^, provide a better correlation with observed mutation rate than steady state transcript levels (**Extended Data Fig. 2a-d**). Using our strand-resolved mutation data, we find that increased transcription decreases the mutation rate for template strand lesions up to 10 nascent transcripts per million (**Fig. 3b**). Beyond this, the mutation rate is not further reduced by additional transcription, suggesting that the remaining mutagenic lesions are largely invisible to TCR (**Extended Data Fig. 2c,d**).

**Fig.3.**
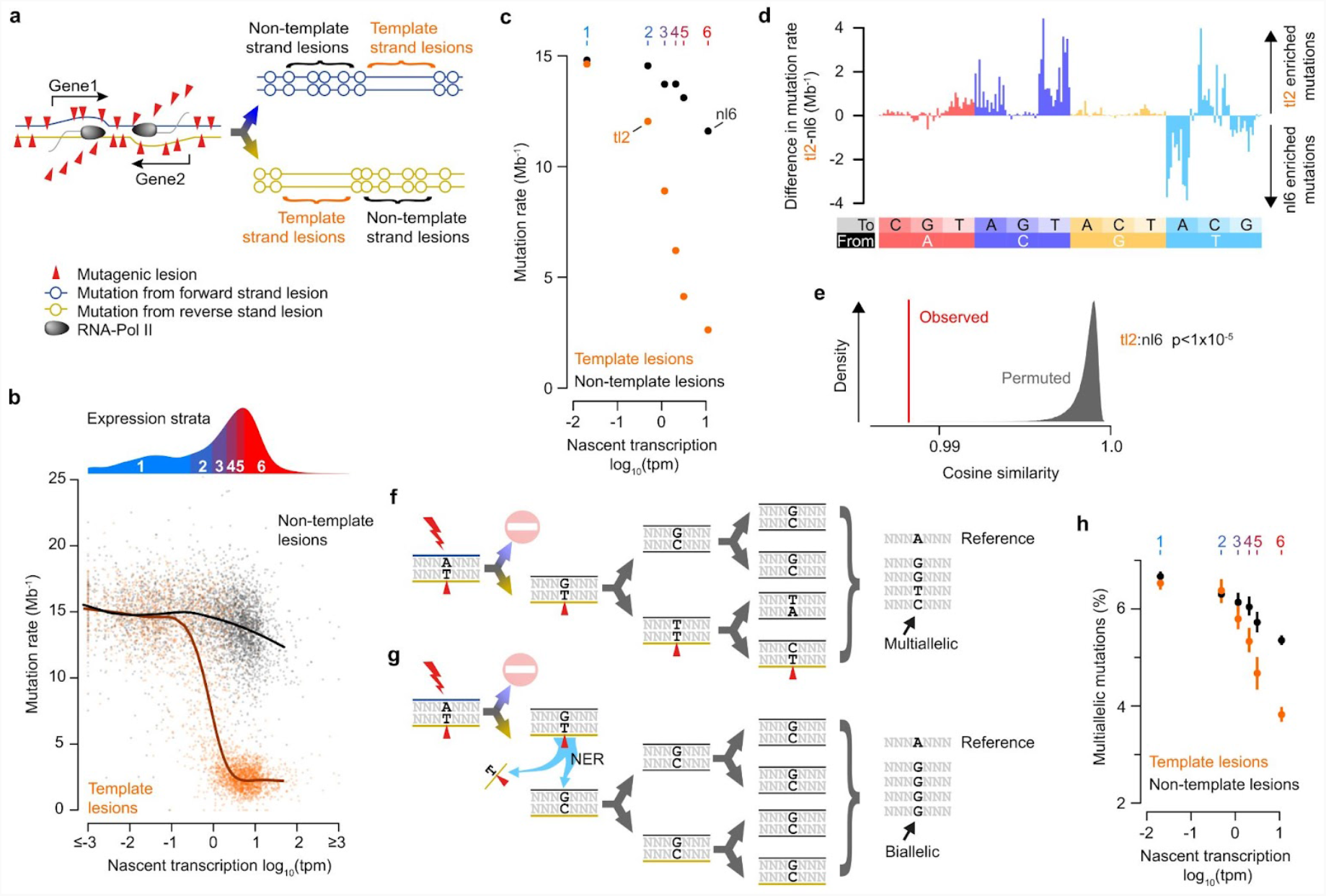
Transcription associated and coupled repair. **a**, DNA lesions (red triangles) on the transcription template strand can cause RNA-polymerase to stall and trigger transcription coupled nucleotide excision repair. Cells that inherit the template strand of active genes are expected to have a depletion of mutations through the gene body. **b**, Mutation rate (y-axis) for individual genes relative to their nascent transcription rate (x-axis) estimated from intronic reads. Mutations rates for each gene (n=3,392) are calculated separately for template (orange) and non-template strand lesions (black); curves show best-fit splines. Grouping of genes into six expression strata, used in subsequent analyses, is indicated by the density distribution above the xy-plot. **c**, Mutation rates for genes grouped into expression strata (1-6 top axis), calculated separately for template strand lesions (orange) and non-template strand lesions (black). 95% bootstrap confidence intervals (whiskers, too small to resolve). Labels indicate data used in subsequent mutation spectra panels (**d**-**e**). **d**, Despite similar mutation rates, the spectrum of mutations differs between non-template lesion stratum 6 (nl6) and template lesion stratum 2 (tl2). **e**, Permutation testing confirms the mutation spectra differs between transcription template and non-template strand, even when overall mutation rates are similar. Comparison of tl2 and nl6 mutation rate spectra (red) and after gene-level permutation of categories, n=10^5^ permutations (grey). **f**, Lesions (red triangles) that persist for multiple cell generations can generate multiallelic variation. **g**, Rapidly repaired lesions persist for fewer cell cycles, and therefore have less opportunity to generate multiallelic variation. **h**, The multiallelic rate (y-axis) for template strand lesions (orange) is reduced with increasing transcription (x-axis). The same is apparent, albeit to a lesser extent, for non-template lesions (black), indicating enhanced repair of non-template lesions is also associated with greater transcription. Whiskers show bootstrap 95% confidence interval.

Non-template strand lesions show a modest reduction in mutation rate with increased transcription (**Fig. 3c**). The signature of TCR on the template strand does not match the profile of reduced mutation rate on the non-template strand of expressed genes, arguing that cryptic antisense transcription is not responsible (**Fig. 3d,e**; **Extended Data Fig. 2e-j**). This indicates that there is either (i) enhanced (non-TCR) surveillance of lesions on the non-template strand or (ii) reduced DEN damage to transcriptionally active genes.

We used another insight from lesion segregation to disentangle patterns of differential damage from differential repair. Since DNA lesions from DEN treatment can persist for multiple cell cycles, each round of replication could incorporate a different incorrectly paired nucleotide opposite a persistent lesion. This results in multiallelic variation: multiple alleles at the same position within a tumour^2^ (**Fig. 3f**). Lesions in efficiently repaired regions will persist for fewer generations and therefore have fewer opportunities to generate multiallelic variation, so are expected to exhibit lower multiallelic rate (the composition-adjusted fraction of mutations with multiallelic variation) than less efficiently repaired regions (**Fig. 3g**). In contrast, differential rates of damage, although influencing overall mutation rate, do not systematically distort the persistence of an individual lesion, so would have no influence on rates of multiallelic variation.

For lesions on the template strand, multiallelic rate decreases with increased transcription (**Fig. 3h**), reflecting the progressive removal of lesions across multiple cell cycles by TCR. The multiallelic rate for non-template strand lesions is also reduced with greater transcription, demonstrating enhanced repair rather than decreased damage. Combined with the distinct repair signature (**Fig. 3d,e**; **Extended Data Fig. 2j**), this indicates a transcription associated repair activity operating in expressed genes that is in addition to the template strand specific TCR. We speculate that this may reflect enhanced global NER surveillance in the more open chromatin of transcriptionally active genes.

### Transcription coupled repair is stochastic

To further explore the mechanisms of TCR, we measured the mutation rate in consecutive 5 kb windows from the transcription start site (TSS) demonstrating subtly (approximately 3.5%) lower mutation rates for both template and non-template strand lesions at the 5’ end of non-expressed genes. This trend was also seen for the non-template strand at all expression strata (**Fig. 4a**; **Extended Data Fig. 3b,c**). We subsequently calculated TCR efficiency as a ratio of the template (observed) versus the corresponding non-template (expected) mutation rates for the same sets of genes, thus negating partial correlates such as 5’ end effects and non-TCR surveillance of expressed genes. We found that TCR efficiency decays slowly: mutation rates increase through the gene body. This finding supports a mechanism in which an RNA polymerase must repair 5’ lesions before downstream 3’ lesions can be repaired.

**Fig.4.**
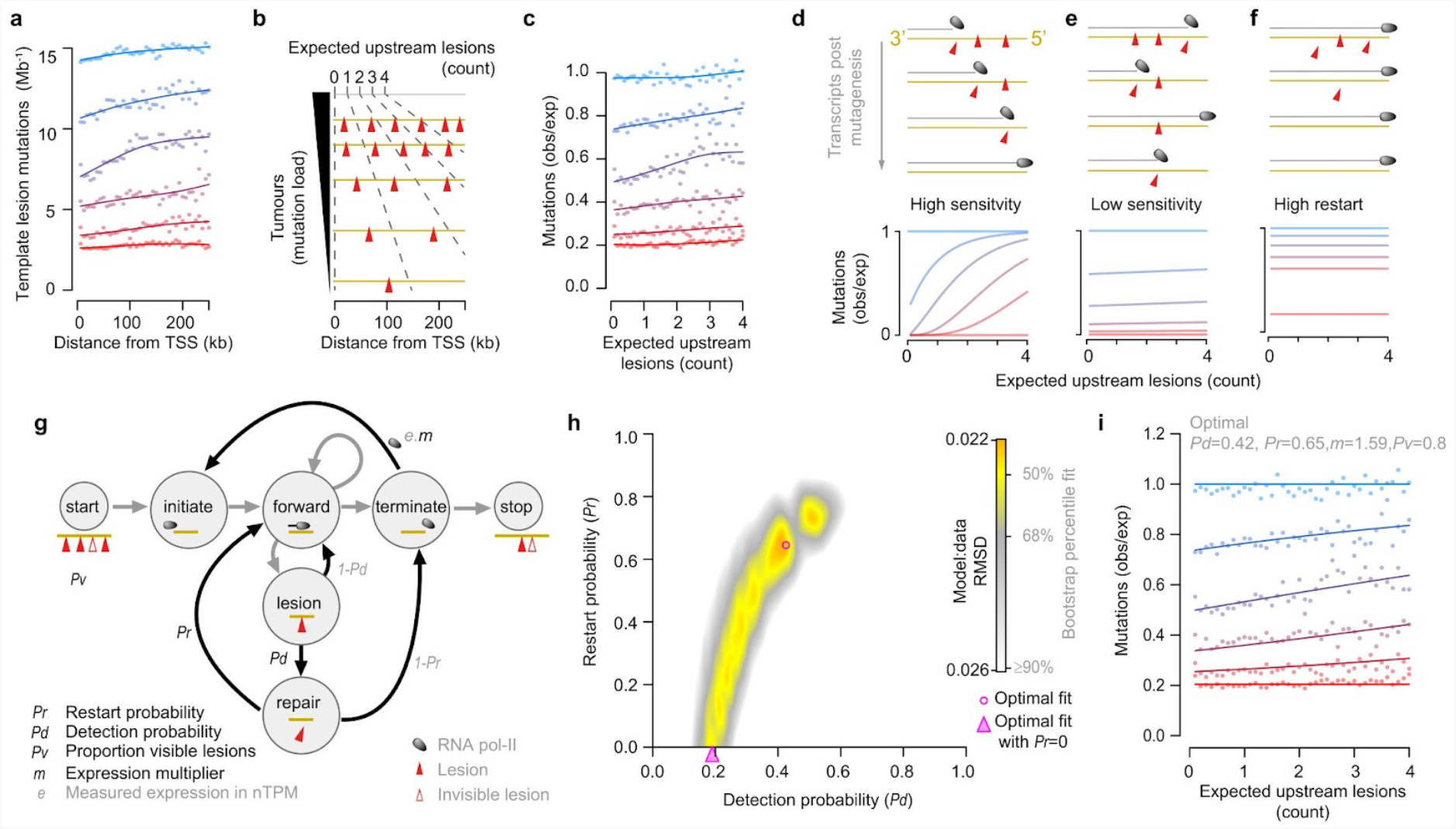
High stochasticity of transcription coupled repair (TCR). **a**, Mutation rates for each expression strata (low to high expression: blue→red, thresholds as per **Fig. 3b**), calculated in 5 kb windows (points) from the TSS for template strand lesions. Curves show best-fit splines. **b**, Schematic of per-tumour normalisation, to calculate the number of expected upstream lesions for each analysis window (Methods). **c**, Genes with intermediate levels of expression (strata 2-5) exhibit a lower mutation rate at their 5’ end. Observed versus expected mutations (y-axis) calculated as the ratio of template to non-template strand. Expected upstream lesion count (x-axis) categories as per **b. d**,**e**,**f**, Alternative models (upper panels) of TCR lesion detection sensitivity and ability to re-start transcription following the triggering of TCR predict different mutation distributions through gene bodies (lower panels). **g**, Mathematical model of TCR dynamics used to generate model profiles in **d**-**f**. A string of nucleotides (yellow line) with DNA lesions (red triangles) is subject to transcription (grey arrows), and probabilistic TCR events (black arrows). On encountering a lesion the probability of its detection (*Pd*) and of polymerase re-start following lesion repair (*Pr*) are independent model variables. The fraction of lesions visible to TCR (*Pv*) and an expression multiplier parameter (*m*) are additional independent variables. Example mutation rate profiles in panels **d**,**e**,**f** were generated by the model using parameters (*Pd, Pr, m, Pv*) of **d** (1,0,1.5,1), **e** (0.2,0.8,5,1), and **f** (0.1,1,1.5,1). **h**, Heatmap showing optimal fits for all grid-search tested values of *Pd* and *Pr* (8.4×10^8^ parameter combinations tested). Optimal fits (pink shapes; circle *Pr*≥0, triangle *Pr*=0) identified from gradient descent exploration initialised by high-quality grid-search fits. Landscape shading from the quantile distribution of fits between the observed data and bootstrap samples of it. **i**, Comparison of optimal model parameters (curves) to observed data (points).

Since the aggregate pattern of mutations is influenced by genic sequence composition and individual tumour mutational burden, we compared observed:expected mutation rates (as above) normalised to the expected number of upstream lesions, rather than genomic distance from the TSS (**Fig. 4b**). This confirms that (i) there is no TCR in the absence of nascent transcription, (ii) TCR efficiency decays approximately linearly with the number of upstream lesions in a gene body, and (iii) highly expressed genes (>10 nTPM) show negligible decay in TCR efficiency through the gene body, indicating that all detectable lesions have been removed (**Fig. 4c**; **Extended Data Fig. 3d-i**).

The linear decay in TCR efficiency through gene bodies is unexpected. If template strand lesions efficiently trigger RNA-polymerase stalling and TCR, then the 5’ end of moderately expressed genes would be cleared of lesions but the 3’ end would remain unrepaired. This would result in a sigmoidal pattern of mutation rates through gene bodies (**Fig. 4d**), which would progressively move towards the 3’ end with increasing transcription rate. Reduced lesion detection sensitivity would flatten the anticipated sigmoidal curve, as would the frequent re-initiation of transcription by RNA-polymerase following the triggering of NER (**Fig. 4e,f**).

To systematically compare these possibilities to the observed mutation data, we defined a mathematical model with four free parameters, exploring plausible values for the probability of lesion detection (*Pd*) and the probability of polymerase restart (*Pr*). The additional model parameters are an expression multiplication factor (*m*) to convert experimental measures of nascent expression to numbers of polymerases, and a measure for the fraction of lesions that are visible to TCR (*Pv*) (**Fig. 4g**). Plausible fits of the observed mutation patterns to this model place an upper bound of 60% lesion detection sensitivity per polymerase traversal, demonstrating that the transcription coupled triggering of TCR is stochastic (**Fig. 4h,i**; **Extended Data Fig. 3j-n**). Model fits that exclude polymerase restart (*Pr*=0) also give essentially optimal (within bootstrap uncertainty) fits to the model; under this scenario the upper bound lesion detection sensitivity is 20%. We conclude that although RNA polymerase is processive, its triggering of NER is highly stochastic, and the resulting high rates of lesion bypass shape the distribution of genic mutations following DNA damage.

### Steric influences on DNA damage and repair

Transcription associated repair of non-template lesions (**Fig. 3h**) implicates DNA accessibility as an important influence on the repair of DNA damage. Accessibility is also influenced by nucleosome positioning and transcription factor binding, both of which have been shown to broadly influence mutation patterns^7,9–11,37^. However, these features are not independent of other correlates of mutation rate, such as gene expression; the relative contributions of DNA damage, replication error, and repair processes remain to be fully resolved. Therefore, we explored how the accessibility of protein-DNA complexes shapes mutagenesis by jointly analysing multiallelic variation and the lesion strand resolved mutation rate.

Within consistently positioned nucleosomes^38^, mutation rates have a 10 bp periodicity (**Fig. 5a**) and higher mutational maxima on the 3’ side of the nucleosome^11,39^. Recapitulation of the mutation rate profile by multiallelic variation rate (**Fig. 5b**) supports the notion that these patterns are shaped by differential repair efficiency rather than differential damage.

**Fig.5.**
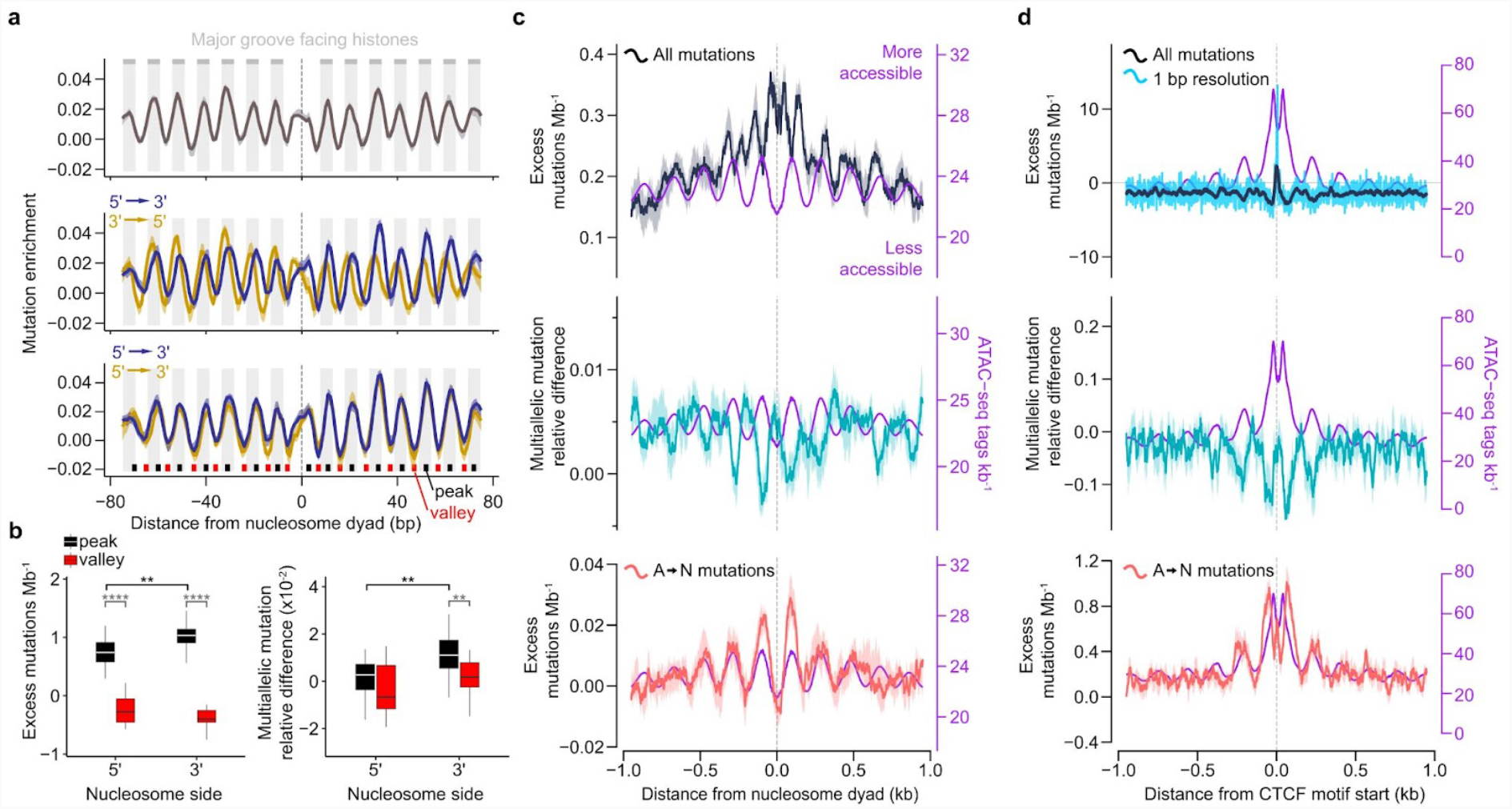
Rapid repair of accessible DNA shapes the mutational landscape but CTCF binding causes extreme local distortions. **a**, The compositionally corrected mutation rate shows helical (10 bp) periodicity over nucleosomes (top). Separating the mutation rates by the lesion containing strand (blue, forward; gold, reverse) reveals two partially offset periodic profiles (middle panel). Orientating both strands 5’→3’ demonstrates that the profiles are mirror images (bottom). Mutation rate peaks (black) correspond to regions where the DNA major groove faces into the histones, and valleys (red) where the major groove faces outward. **b**, For the lesion containing strand, mutation rates are significantly higher for the peaks on the 3’ side of the nucleosome dyad than on the 5’ side (** p=0.0012, two tailed Wilcoxon test). Comparing the compositionally corrected multiallelic rates, shows significantly increased multiallelic variation for the 3’ peaks (** p=0.0026, two tailed Wilcoxon test), indicating the increased mutation rate results from slower repair on the 3’ side of the dyad. **c**, At a broader scale, nucleosome occupancy also shapes the mutational landscape, with higher mutation rates (top panel, black) over the nucleosomes (x=0 and low accessibility as measured by ATAC-seq, purple) and lower rates in the linker regions (ATAC-seq peaks). ATAC-seq from P15 mouse liver. High rates of multiallelic variation (green, mid panel) are found at sites of low accessibility and high mutation rate, indicating that high rates of mutation represent slow repair. The rate of A→N mutations is the inverse of the overall mutation profile (pink, lower panel), with high rates of A→N corresponding to accessible regions and rapid repair. **d**, Mutation rates are dramatically elevated at CTCF binding sites (black, 21 bp sliding window; blue single base resolution). Panels as described for c. High accessibility (purple) again corresponds to low multiallelic variation (green) and low mutation rates. Mutations of A→N (pink) closely track DNA accessibility. The increase in mutation rate within the binding site (top panel, blue) is not accompanied by a similar increase in multiallelic rate indicating it cannot be explained by suppressed repair.

In highly accessible regions between nucleosomes there are low rates of mutations and also low rates of multiallelic variation (**Fig. 5c**), indicating rapid removal of lesions in these linker regions; this pattern was largely consistent for strand phased mutations from T, C, and G lesions. Unexpectedly, the opposite was found for mutations apparently arising from A lesions, with high rates of A→N in accessible regions (**Fig. 5c**; **Extended Data Fig. 4a-d**).

As with nucleosomes, the binding of CTCF influences mutation rates in cancer^37,40^. At experimentally determined CTCF binding sites (in P15 mouse liver, Methods) we found a pronounced hotspot of mutations, but without the corresponding increase in multiallelic rate expected from suppressed repair (**Fig. 5d**). This implicates the local elevation of DNA damage as the source of the mutation hotspot. In contrast, accessible regions (identified in P15 mouse liver ATAC-seq, Methods) adjacent to CTCF binding sites have low mutation rates and correspondingly low multiallelic variation rates, indicating more efficient repair (**Fig. 5d**).

At single nucleotide resolution (**Fig. 6a**), the net enrichment of mutations at CTCF binding sites is highly strand-and position-specific. In general, the mutation rate is elevated at nucleotide positions where the CTCF protein contacts neither, or only one, strand of the DNA backbone; these bases and other backbone are solvent exposed, but the DNA helix remains occluded by CTCF (**Fig. 6a**, motif nucleotide positions -4 to 3). A notable outlier is position 6 of the binding motif, which has close CTCF contacts on the base and backbone of both strands and yet exhibits a strong T→N mutation enrichment for the motif C-rich strand.

**Fig.6.**
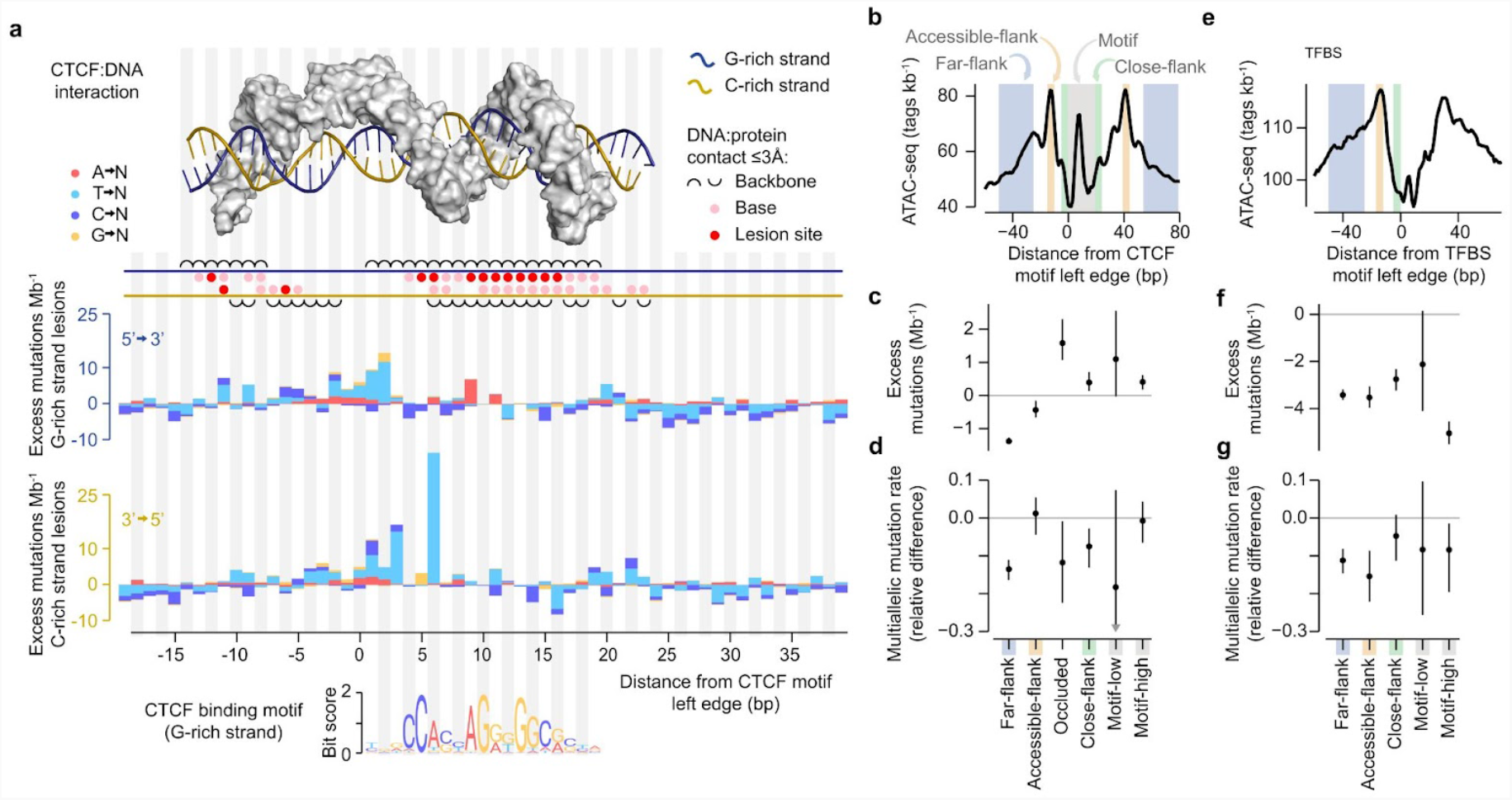
Mutation enrichment and depletion at transcription factor binding sites (TFBS). **a**, The molecular structure of the CTCF:DNA interface (top) reflects the strand specific mutation profiles of CTCF binding sites (histograms, composition corrected). A composite crystal structure of CTCF zinc fingers 2-11 (grey surface) is shown binding DNA (blue & gold strands) and close protein:DNA contacts (≤3Å) illustrated below the structure. At nucleotide positions with close contact between CTCF and atoms thought to acquire mutagenic lesions (red circles), the corresponding strand specific mutation rates are generally lower than genome-wide expectation (y≤0; excepting apparent A→N mutations considered later). Mutation rates are high (y>0) for nucleotide positions with backbone-only contacts or no close contacts but still occluded by CTCF. CTCF motif position 6 exhibits an exceptionally high T→N mutation rate that cannot be readily reconciled with the structure, but the strand specificity demonstrates it is a consequence of DEN exposure. **b**, The profile of DNA accessibility around CTCF binding sites, defines categories of sequence (shaded areas) considered subsequently. **c**, Mutation rates are higher than genome-wide expectation (y=0) for CTCF binding motif nucleotides and their close flanks. **d**, This is not reflected in increased rates of multiallelic variation. CTCF occluded positions (positions -5 to 3 of the CTCF motif) show the greatest elevation of mutation rate but evidence of decreased multiallelic variation. Both high information content (motif-high, bit score>0.2) and low information content (motif-low, bit-score ≤0.2) motif positions have high mutation rates. **e**, DNA accessibility around non-CTCF transcription factor binding sites (TFBS) aligned as in **b. f**,**g**, In contrast to the situation for CTCF, all TFBS categories of sites have suppressed mutation rate compared to genome-wide expectation (**f**, y=0), and suppression of multiallelic variation (**g**) indicates enhanced repair. However, high information content motif sites (motif-high) have exceptionally reduced mutation rate not similarly reflected by multiallelic variation, suggesting there may be reduced damage in addition to efficient repair at these sites.

Despite the localised increases in mutation rates within the CTCF binding footprint, multiallelic rates are not increased, and for some categories of sequence are modestly decreased compared to genome wide expectation (**Fig. 6b-d**). This suggests that the elevated mutation is not primarily a consequence of suppressed repair, rather enhanced DNA damage within CTCF binding sites. The pattern of enhanced damage appears specific to CTCF and does not generalise to other liver-expressed transcription factors (**Fig. 6e-g**) which exhibit reduced mutation rates and multiallelic variation both adjacent to and within their binding sites, implicating more efficient repair of lesions within TFBS.

High information content binding site nucleotides within TFBS show exceptionally reduced mutation rates, even compared to low information content nucleotides within the motifs. The reduced mutation rate is not reflected by more pronounced reductions in multiallelic variation (**Fig. 6f,g**). This raises the possibility that close contacts between a binding protein and atoms susceptible to mutagen attack could offer some protection from lesion formation.

The anomalous enrichment of apparent A→N mutations on the lesion containing strand is a recurrent feature of genomic loci with high efficiency repair, including linker regions between nucleosomes, and adjacent to CTCF and transcription factor binding sites (**Fig. 5c,d**; **Fig. 6**). The enrichment of A→N mutations also extends into sequence specific binding sites (**Fig. 6a**; **Extended Data Fig. 4e,f**). A possible explanation for the specific enrichment of A→N mutations is that, in some circumstances, the activity of NER is itself mutagenic.

### Nucleotide excision repair is mutagenic

We propose a mechanistic model for mutagenic NER, arising when two lesions occur in close proximity, but on opposite strands of the DNA duplex. Repair of one lesion, which entails excision of a ∼26 nt single stranded segment containing the lesion^41,42^, would leave a single stranded gap containing the second lesion on the opposite strand; resynthesis using this as a template would necessitate replication over that remaining lesion (**Fig. 7a**). As a result, nucleotide misincorporation opposite a T lesion in the single stranded gap would be erroneously interpreted as a mutation from an A lesion (**Fig. 7a**) when phasing lesion segregation. We subsequently refer to this mechanism as translesion resynthesis induced mutagenesis (TRIM), or NER-TRIM specifically in the context of NER.

**Fig.7.**
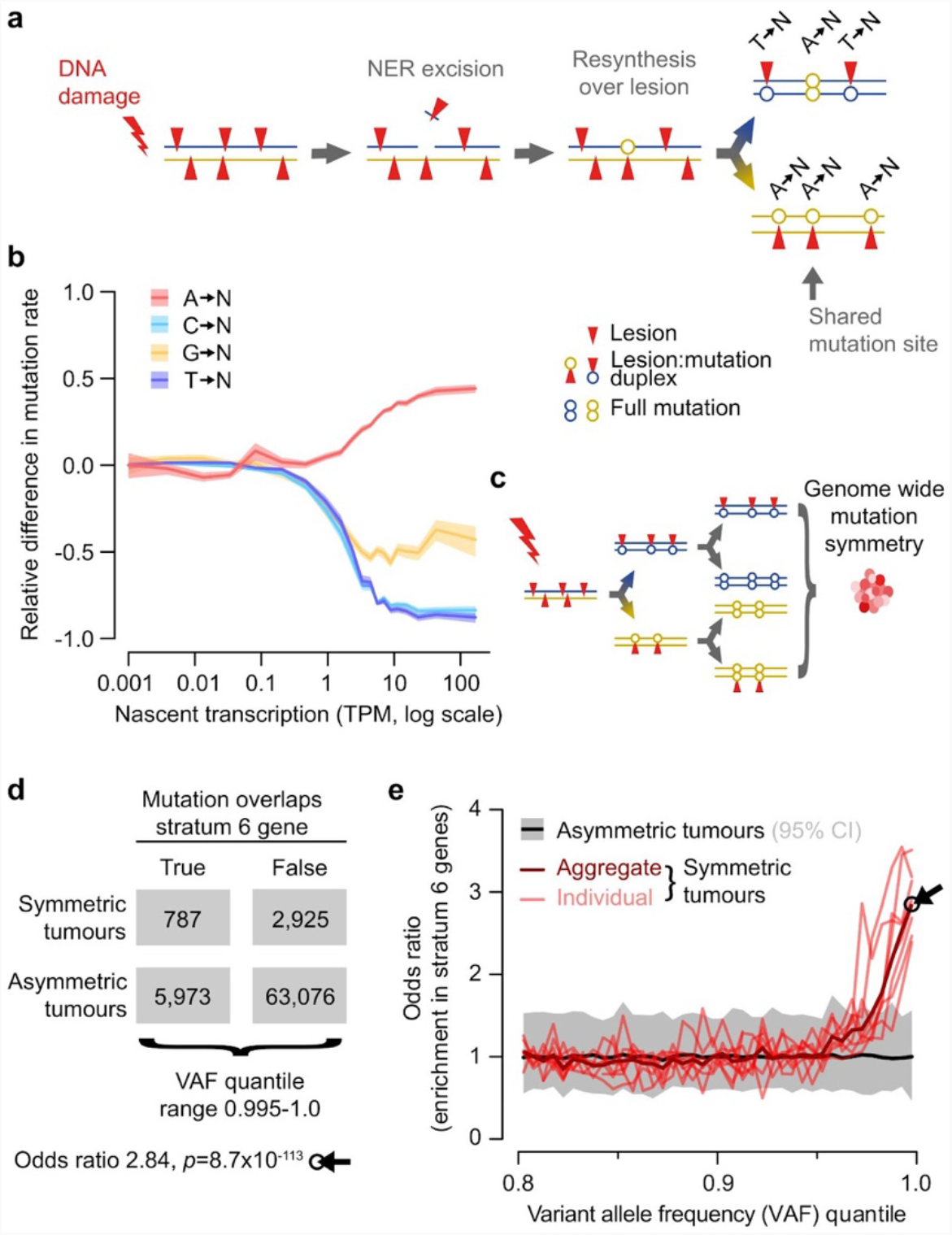
Nucleotide excision repair (NER) is mutagenic when lesions on opposing strands are in close proximity. **a**, Mechanism of NER translesion resynthesis induced mutagenesis (NER-TRIM). A lesion-containing single stranded stretch of DNA is excised and consequently a residual lesion on the opposite strand is used as a low fidelity template for the repair synthesis. This creates isolated mutations with opposite strand asymmetry to the genomic locality (e.g. A→N within a T→N segment). Most lesion induced mutations are not shared between the two daughter lineages, whereas those from NER-TRIM can be shared (grey arrow). **b**, The rate of A→N mutations on the genic template strand increases with gene expression, mirroring the decrease in mutations from other bases, due to transcription coupled repair. The relative difference (y-axis) of mutation rate for each nucleotide is (obs-exp)/(obs+exp); exp is the mutation rate for that nucleotide in non-expressed genes (nascent TPM <0.001), and obs is the rate observed in the body of genes with the indicated expression level (x-axis). Rates shown for lesions on the transcription template strand, 95% C.I. (shaded areas) from 100 bootstrap samples of genes. **c**, Schematic illustrating the generation of a mutationally symmetric tumour through the survival of both post-mutagenesis daughter genomes. Such symmetric tumours are expected to contain NER induced mutations characterised by abnormally high VAF, because they will be shared by both contributing genomes (**Extended Data Fig. 5b**). **d**, Contingency table illustrating the enrichment of mutations with high VAF (0.995-1.0 quantile) in highly expressed genes of mutationally symmetric tumours (n=8) compared to mutationally asymmetric tumours (n=237). Statistical significance by two-tailed Fisher’s exact test. **e**, Symmetric tumours are highly enriched for high VAF mutations in highly expressed genes. Odds ratios (y-axis) as in **d**, for VAF quantile bins of 0.005 (x-axis). Black arrow shows odds ratio calculated in **d**.

Since NER-TRIM requires lesions on both DNA strands, mutagenic NER can only occur when both lesion-containing strands are duplexed, for example in the first cell generation following DEN mutagenesis. NER-TRIM cannot occur in daughter cells with only one lesion containing strand. Consequently, regions with the highest -and thus fastest - repair rates are most likely to experience NER-TRIM. This prediction is consistent with our observation of local enrichment of apparent A-lesion mutations in accessible regions with otherwise low rates of mutations and low multiallelic variation (**Fig. 5c**).

Local gradients in repair efficiency are also expected to lead to enrichment of NER-TRIM. The most efficient repair we observed is transcription coupled NER, in which there is a strong gradient of repair efficiency between the template and non-template strands. There is a pronounced increase in the rate of apparent A→N mutations on the template strand of expressed genes, whose sigmoidal profile closely mirrors the decrease in T→N mutations on the same strand (**Fig. 7b**). Importantly, the saturation of repair at higher expression levels is reflected in a corresponding saturation of NER-TRIM, demonstrating that the rate of template strand A→N mutations is not simply dependent on transcription, but on transcription coupled repair.

Similar local gradients of repair can also explain the elevated rate of A→N mutations in CTCF and transcription factor binding sites (**Fig. 6**; **Extended Data Fig. 4e,f**), where there is a gradient between more accessible nucleotides adjacent to the binding site and those nucleotides within the binding site. In this scenario, high efficiency repair of the accessible DNA would result in an excision gap that extends into the binding site, where a more protected lesion then serves as a template for repair resynthesis.

A characteristic prediction of the NER-TRIM model is that both daughter lineages of the originally mutagenised cell can contain a mutation at the same site after NER-TRIM (**Fig. 7a**). In contrast, other DNA damage induced mutations would not be shared between daughters (**Fig. 1a**). A subset of tumours in our dataset provide an opportunity to directly test this mutation-sharing prediction. Of the original DEN induced tumours^2^, 2% (n=8) exhibited the same mutation spectra as other tumours but completely lacked the mutational asymmetry of lesion segregation (**Extended Data Fig. 5a**). This pattern is expected to result from the persistence of mutations derived from lesions on both strands (**Fig. 7c**; **Extended Data Fig. 5b**). Based on genomic and histological evidence, we conclude that these eight mutationally symmetrical tumours are each made up of two diploid sister clones (**Extended Data Fig. 5c-h**). Since these tumours contain both daughter lineages of the initial mutagenised cell, they provide an opportunity to test the mutation sharing expectations of both lesion segregation and NER-TRIM.

The variant allele frequency (VAF) of a somatic mutation is proportional to the fraction of cells in the tumour that contain the mutation. Consequently, we expect the VAF of shared mutations derived from NER-TRIM to be approximately twice that of mutations found in only one of the two daughter cell lineages. Due to the absence of mutational asymmetry in these eight tumours, it is not possible to define which individual mutations arose from NER-TRIM. However, since we have shown that NER-TRIM is enriched in highly expressed genes, we tested if high VAF mutations were biassed to those regions in the symmetrical tumours (n=8) compared to the asymmetric tumours (n=237). Our results demonstrated a pronounced and significant enrichment as predicted, both in aggregate (Odds ratio 2.84, two tailed Fisher’s test p=8.7×10^−113^; **Fig. 7d**) and individually for each tumour (**Fig. 7e**), confirming expectations of the NER-TRIM model.

Finally, we note that in the symmetrical sister-clone tumours, the oncogenic driver mutations in the MAPK pathway that typify these DEN induced tumours^2,21^ are all significantly biassed to the highest VAF mutations, in contrast to the VAF of driver mutations in the asymmetric tumours (p=3.61×10^−5^ two tailed Wilcoxon rank sum test, Bonferroni corrected; **Extended Data Fig. 5i-y**). This suggests that driver mutations in the symmetrical tumours arose through NER-TRIM, and may explain the coevolution of both sister clones in a single tumour.

## Discussion

In damaged DNA, most mutations arise from replication bypass of unrepaired lesions, which can result in chromosome-scale mutational asymmetry^2^. We leveraged this discovery to explore the mechanisms of mutagenesis and repair *in vivo* at high resolution, with single-base, single-strand specificity. By combining the rate of multiallelic mutations, caused by the persistence of DNA lesions for multiple cell generations, with mutation rates we were able to discriminate the relative contributions of initial DNA damage from subsequent repair.

Despite the enzymatic and mechanistic differences in leading and lagging strand replication^7,27–29^ their fidelity appears symmetrical on undamaged DNA^43^. It has long been argued that damaged DNA templates break this symmetrical replication fidelity^23,24,44^. Our results substantially revise this paradigm. In contrast to observations for bulky DNA adducts such as pyrimidine dimers^5,8^, we find symmetric replication fidelity is preserved for smaller adducts, including base ethylation from DEN^20^. Matched patterns of collateral mutagenesis point to recruitment of identical translesion polymerases for on-the-fly bypass of small adducts, on both the leading and lagging strands, thus preserving the mutational symmetry of replication.

Our results demonstrate that, as for replication, lesion bypass of small mutagenic adducts is also a common feature of transcription. Modelling the decay of transcription coupled repair efficiency through gene bodies, indicates that RNA polymerase triggering of NER is highly stochastic and means that small unrepaired lesions are unlikely to be an efficient barrier to gene expression. This observation, especially the presence of 5’ repair bias, is important when there are multiple lesions per gene, and has implications for both accurate modelling of mutation patterns and prediction of oncogenic selection^45^.

In addition to the expected transcription coupled repair of lesions on the template strand, we also used multiallelic variation rates to demonstrate transcription associated repair of non-template strand lesions. In prokaryotes, this phenomenon can occur as a consequence of antisense transcription^46^, but mutation spectra reveal that is not the case in these mammalian tumours. We therefore propose that transcription associated repair represents enhanced genome-wide NER surveillance of more accessible, expressed genes.

Beyond the effects of transcription, the mutational landscape of genomes closely tracks DNA accessibility. This pattern is mirrored by the rate of multiallelic variation, thus providing *in vivo* evidence that more efficient repair of accessible DNA, rather differential DNA damage, is primarily responsible for shaping the distribution of damage induced mutations. Despite multiple reports of elevated mutation rates at transcription factor binding sites^37,40,47,48^, we observed locally reduced mutation rates within, and adjacent to, transcription factor binding sites. This reduction was recapitulated by multiallelic rates, indicating that the binding of transcription factors to DNA does not generally impede efficient repair, and reflects the dynamic nature of such protein-DNA interactions^48,49^.

A notable exception to this occurs at CTCF binding sites which, like stably bound Reb1 and Abf1 in yeast^50^, generate damage hotspots. Specific nucleotides within the CTCF binding footprint show exceptionally elevated mutation rates, and the absence of increased multiallelic variation at these sites implicates enhanced damage rather than exceptionally suppressed repair. The identity of sites with elevated mutation can be only partially reconciled with the structure of the CTCF:DNA interface. We speculate that this structure may be modified, for example by interacting with cohesin, leading to bending^51,52^ and partial melting of the DNA duplex, resulting in greater exposure of the nucleotide bases to chemical attack.

Finally, we found that genomic regions that are most efficiently repaired are also, counterintuitively, specifically prone to repair induced mutagenesis. This mechanism of NER template resynthesis induced mutagenesis (NER-TRIM) results from the repair of lesions in close proximity, but on opposite strands. It is therefore expected to occur when damage loads are high or closely spaced, for example UV damage in promoters and ETS factor binding sites^48,53^. While NER-TRIM mutations represent only a small fraction of damage induced mutations, they are specifically biassed to functionally important sites, and are responsible for most driver mutations seen in symmetric tumours. Perhaps most importantly, NER-TRIM preferentially results in the misincorporation of a normal DNA base on the template strand of highly expressed genes. That incorrect normal base is not a substrate for subsequent repair and could therefore lead to efficient mis-coding of a protein before genome replication. In the case of an oncogenic mutation, potentially driving otherwise quiescent cells towards oncogenic transformation.

Our ability to resolve both mutation rates and multiallelism at single-strand, single-base resolution allows us to infer lesion longevity and thus disentangle differential DNA damage from differential repair. This powerful approach provides *in vivo* insights into how strand-asymmetric mechanisms underlie the formation, tolerance, and repair of DNA damage, thereby shaping cancer genome evolution.

## Methods

### Genomic annotation

The C3H/HeJ mouse strain reference genome assembly C3H_HeJ_v1^54^ was used for read mapping, annotation, and analysis. WGS regions with abnormal read coverage (ARC regions, 12.7% of the genome) were masked from analysis, as previously described ^2^. Gene annotation was obtained from Ensembl v.91^55^.

### Mutation asymmetry

Mutation calling and quality filtering, was performed using WGS of 371 DEN induced liver tumours from n=104 male C3H mice, as previously reported^2^. All mutation data was derived from sequence data in the European Nucleotide Archive (ENA) under accession PRJEB37808 and processed files directly used as input for this work are publicly available https://doi.org/10.1038/s41586-020-2435-1.

Genomic segmentation on mutational asymmetry was performed as previously reported^2^. Mutational strand asymmetry was scored for each genomic segment using the relative difference metric *S=(F-R)/(F+R)* where *F* is the rate of mutations from T on the forward (plus) strand of the reference genome and *R* the rate of mutations from T on the minus strand (mutations from A on the plus strand). A mutational asymmetry score of *S* >0.33 was used to identify the inheritance of forward strand lesions and *S* <-0.33 as the inheritance of reverse strand lesions. A rare subset of tumours (2.7%) exhibited uniform mutational symmetry (>99% of autosomal mutations in genomic segments with *abs(S)* < 0.2, these were labelled “symmetric” tumours.

Except where otherwise stated (within first and final results sections), analyses were confined to n=237, clonally distinct DEN induced tumours that met the combined criteria of: (i) not labelled as symmetric, (ii) tumour cellularity >50%, and (iii) >80% of substitution mutations attributed to the DEN1 signature by sigFit (v.2.0)^56^.

Relative to the reference genome sequence, a plus (*P*) strand gene is transcribed using the reverse (*R*) strand as a template. So a *P* strand gene in a genomic segment with *R* strand lesions (denoted *RP* orientation) is expected to be subject to transcription coupled repair. A minus strand (*M*) gene with forward (*F*) strand lesions (*FM* orientation) is also expected to be subject to transcription coupled repair, as the retained lesions are again on the transcription template strand. Conversely *FP* and *RM* orientation combinations will have lesions on the non-template strand for transcription. For DNA replication, we similarly refer to whether the preferential template for the leading strand contains the retained lesions or whether the preferential template for the lagging strand contains the retained lesions.

### Mutation rates and spectra

Mutation rates were calculated as 192 category vectors representing every possible single-nucleotide substitution conditioned on the identity of both the upstream and downstream nucleotides. Each rate being the observed count of a mutation category divided by the count of the trinucleotide context in the analysed sequence. To report a single aggregate mutation rate, the three rates for each trinucleotide context were summed to give a 64 category vector and the weighted mean of that vector reported as the mutation rate. The vector of weights being the fraction of each trinucleotide in a reference sequence, for example the composition of the whole genome. Strand-specific mutation rates were calculated with respect to the lesion containing strand, with both mutation calls and sequence composition reverse complemented for reverse strand lesions. Autosomal chromosomes were considered diploid and the X chromosome haploid (male mice) for the purposes of calculating mutation rates and sequence composition. For the counting of strand-specific mutations, a threshold variant allele frequency (VAF) >10% was applied to remove mutation calls from contaminating non-clonal cells.

Subtracted spectra plots (**Fig. 2c,d**) were calculated by subtracting the counts of simulated tumour datasets from those of observed datasets and then scaling as for mutation spectra so the absolute area of the histogram sums to 100. Percent repair efficiency (**Extended Data Fig. 2j**) was calculated as *(observed/expected) × 100* where expected was the corresponding mutation rate for non-expressed genes (stratum 1, see below) averaged between the template and non-template strand. Cosine similarity was used as a relative measure of mutation signature similarity. Mutation signature deconvolution was performed using sigFit (v.2.0), with two component signatures (*K*=2) chosen based on heuristic goodness-of-fit for integer values of *K* from 2 to 8, with 2,000 iterations each. Final *K*=2 deconvolution used 40,000 iterations.

The expected number of mutations at each position of the analysed TFBS and nucleosome regions was calculated as a sum of genome-wide rates (mutations bp^-1^) for that particular trinucleotide context from each tumour that had this region classified as either forward or reverse segment. The genome-wide rate for each tumour was calculated by dividing the number of mutations in a particular trinucleotide context (that fall within genomic space phased to have inherited either forward or reverse lesion-containing strand) by the total count of that trinucleotide in that genomic space; this was done separately for forward and reverse segments.

Excess mutations Mb^-1^ were calculated as *(observed*_*i,n*_*-expected*_*i,n*_*)×10*^*6*^*/(count*_*i*_), where *i* is the relative position within the region, *count*_*i*_ represents a total number of regions with non-’N’ nucleotide at position *i*, and *n* is specific mutation context (e.g. mutation from A). Mutation enrichment was calculated as *(observed*_*i,n*_*-expected*_*i,n*_*)/(observed*_*i,n*_*+expected*_*i,n*_). Rolling mean values were plotted using windows of 51 bp and 21 bp for nucleosome and CTCF centred plots, respectively. 95% confidence intervals were calculated based on bootstrap sampling of the analysed regions.

### Multiallelic mutation rates

Aligned reads spanning genomic positions of somatic mutations were re-genotyped using Samtools mpileup (v.1.9)^57^. Genotypes supported by ≥2 reads with a nucleotide quality score of ≥20 were reported, considering sites with two alleles as biallelic, those with three or four alleles as multiallelic. For a defined set of mutations, the background composition is the count of mutations in each of the 64 possible trinucleotide contexts. The count of multiallelic mutations in each of those 64 categories is divided by the corresponding background mutation count and the weighted average of those ratios reported as the multiallelic rate. As for mutation rates, the vector of weights being the fraction of each trinucleotide in a reference sequence, for example the composition of the whole genome.

### Gene expression

Paired-end, stranded total RNA-seq from unexposed P15 C3H male mouse livers (n=4, matching the developmental time of mutagenesis) were aligned, annotated, and quantified as previously reported^2^. All transcriptome data used was derived from sequence data in Array Express under accession E-MTAB-8518 and is publicly available https://doi.org/10.1038/s41586-020-2435-1.

The transcription strand of RNA-seq reads was resolved using read-end and mapping orientation using Samtools (v.1.7.0) and read-pairs exclusively mapping within annotated exons identified using Bedtools intersect (v.2.29.2)^58^. Intronic read-pairs were defined as those mapping within a genic span, derived from a sense-strand transcript and not in the exonic set.

For genes with multiple annotated transcript isoforms, the sum of transcripts per million (TPM) over the isoforms was taken as the expression measure (mature transcript, steady-state), though similar results with the same conclusions were obtained if the maximum for any one isoform was used. Nascent transcription was quantified by counting read-pairs with a mapping quality (MAPQ) of >10 overlapping intronic regions (defined as intronic in all annotated transcript isoforms of the gene) using Bedtools multicov (v.2.29.2). The read count was normalised to reads per kilobase of analysed intron for each gene in each sequence library, and then normalised to transcripts per million (TPM) for each library. The final nascent transcript expression estimate per gene was taken as the mean of nascent TPM over replicate libraries. Nascent transcription estimates could be generated for 85% (n=17,304) of protein coding genes.

Gene-based analyses of mutation rates used the genomic extent of the most highly expressed transcript isoform (the primary transcript) based on P15 C3H mouse liver gene expression. Overlapping genes, defined by primary transcript coordinates, were hierarchically excluded from analysis. Starting with the most expressed gene, any overlapping less-expressed genes were excluded. For the plotting of per-gene, per-strand mutation rates (**Fig. 3b**; **Extended Data Fig. 2b-d**) only genes spanning >2 million nucleotides of strand resolved tumour genome aggregate were shown (n=3,392 genes) to minimise stochastic noise from genes with little power individually to accurately estimate mutation rates. Analyses aggregating rates by expression bin included all genes within the bin.

Genes with similar estimates of nascent expression were aggregated for analysis of transcription coupled repair. The sigmoidal distribution relating nascent transcription rate to mutation rate (**Fig. 3b**) was segmented using linear regression models in the R package Segmented (v.1.3-3)^59^. This defined n=4,649 genes with zero or low detected nascent expression (<0.287 TPM) in which reduced mutation rates associated with transcription coupled repair are essentially undetectable; subsequently stratum 1 genes (light blue in plots). Genes expressed at a greater rate than segmentation threshold (>3.73 TPM) do not show a further decrease in mutation rate with increased expression; these n=7,176 highly expressed genes were defined as stratum 6 (bright red in plots). The n=4,005 genes with intermediate expression (0.287-3.73 TPM) exhibited a log-linear relationship between expression and mutation rate. These were quantile split into strata 2 to 5, containing approximately 1,000 genes in each strata.

### Replication strand bias

Replication fork directionality (RFD) is a relative difference metric that scales from 1 to -1. RFD values >0 indicate a consensus rightward progressing replication fork, whereas RFD <0 indicates a consensus leftward progressing fork. RFD values from Okazaki fragment sequencing (OK-seq) of mouse primary splenic B cells^60^ were obtained as BED files calculated in 1 kb consecutive windows. Experimental replicates were combined by taking mean RFD values for each window. The original BED coordinates corresponding to GRCm38 reference genome coordinates were projected into the C3H_HeJ_v1 reference assembly using halLiftover (v.2.1) ^61^ through murine whole genome alignments^54^. Windows reduced to 500 bp or less were removed from the analysis.

For each DEN induced tumour we identified all RFD segments that were completely contained within lesion segregation mutational asymmetry segments (as defined above) with abs(*S*) >0.33. For these segments we resolved the lesion containing strand to the template of either the leading or lagging replication strand. A forward strand mutation asymmetry (lesions on the forward strand, *S* >0.33) and rightward progressing replication fork (RFD >0) is consensus lagging strand replication over the lesions (**Fig. 1e**). Similarly *S* <-0.33 and RFD <0 is also lagging strand replication over lesions. Consensus leading strand replication over lesions is indicated by *S* >0.33, RFD <0; or *S* <-0.33, RFD >0.

RFD values were attributed to bins ranging from -1 to 1 in increments of 0.1. Genic and intergenic windows with RFD values were derived using bedtools intersect and subtract against protein coding genes in the C3H gene annotation, respectively. Bootstrap sampling (n=520) was conducted for a given sample by randomly drawing windows with RFD measures with replacement, at a frequency matching the number of unique windows occurring across phased segments.

### Mutation clusters

For each nucleotide substitution mutation, the closest adjacent mutation was found. Null expectations of mutation spacing were generated by sampling mutation positions from other tumours without replacement, to generate an identical number of proxy mutations for each tumour. Initial analysis of mutation spacing indicated strong enrichment of mutations spaced <11 nt apart and evidence of enrichment to 100 nt spacing. Mutation clusters were defined as chains of mutations within the same tumour spaced <*X* nt from adjacent mutations, with *X*=11, *X*=101, or *X*=201 depending on analysis as indicated. Over 97% of *X*=101 mutation clusters (29,307/30,028) contained only two mutations, 721 clusters contained three mutations, and no larger clusters were identified. 100% of *X*=101 clusters from proxy-tumour mutations contained only two mutations.

For each mutation cluster, if it was located within a lesion segregation mutation asymmetry segment, we annotated the mutations within the cluster with respect to the inferred lesion containing strand. For a genomic segment containing reverse-strand lesions, the leftmost mutation site would be the first used as a template for an extending DNA polymerase (as DNA synthesis extends 5’→3’), and the rightmost mutation site replicated over subsequently.

These orientations are reversed for a genomic segment containing forward-strand lesions. The first replicated-over mutation site for each cluster was annotated distinctly from subsequent sites in the cluster.

Pairs of mutations were phased to the same chromosome by co-occurrence in the same sequencing read. Sequencing reads were extracted from genomic alignments using Samtools mpileup (v.1.7) where they overlapped both genomic positions of a pair of mutations called from the same tumour and separated by ≤75 nt. Any sequencing read supporting the called mutant allele with a phred-scaled quality score ≥20 at both mutation positions was taken as support for those mutations occurring on the same chromosome.

Mutation clusters were resolved to preferential leading or lagging strand replication based on both RFD ^60^ and *S* as described above, such that clusters annotated as lagging strand synthesis would use a lesion containing template strand for lagging strand synthesis. Only the more extreme RFD windows (|RFD| >0.25) were considered for comparisons of leading versus lagging strand asymmetry, so that any strand differences were not swamped by regions with low levels of replicative asymmetry. Clusters were defined with *X*=101 as above resulting in n=5,375 leading strand and n=6,262 lagging strand clusters, the difference in count attributable to transcription coupled repair correlating with leading strand replication (**Fig. 1f**). Accounting for base substitution mutation rate differences there is no significant difference in mutation cluster rate between leading and lagging strands (Two tailed Fisher’s test odds ratio 1.0, *p*=0.8). Cluster length distributions were compared using a two-sample, two-sided Kolmogorov-Smirnov test (ks.test function in R). To estimate statistical power for detecting differences in cluster size distribution between leading and lagging strands, we simulated distorted length distributions. The lagging strand length distribution vector was partitioned into clusters of length ≤10 (short) or >10 (long) and randomly sampled with replacement to produce a vector of length matching the leading strand vector. Bias sampling between the short and long cluster bins was controlled by parameter *d*. An undistorted sample of the original distribution would be *d*=0; whereas 10% of short clusters sampled from the long bin instead of the short bin would be *d*=0.1. Two-sample, two-sided Kolmogorov-Smirnov tests comparing the original to the distorted sample distribution were applied to 100 bootstraps for each tested value of *d* (0 to 0.1 in increments of 0.0005), recording nominal significant difference at p<0.05. The percent of bootstraps supporting nominal significance is the power to detect significance at the tested value of *d*.

### Transcription coupled repair

Annotated genes (Ensembl v.91) were partitioned into six expression strata based on P15 liver RNA-seq (see above). For each tumour, genes were identified that were wholly contained within a mutation asymmetry segment. Using the annotated transcriptional orientation of the gene and mutational asymmetry of the tumour, each of these genes was categorised as either template strand lesion or non-template strand lesion.

Considering TCR efficiency through gene bodies, starting at the annotated 5’ end (transcription start site, TSS), the genomic span of each gene was partitioned into consecutive 5 kb segments. Final segments of less than 5 kb span were discarded. The mutation rate was calculated using the 192 category unfolded mutation spectra (see above) for the *i*^th^ 5 kb window position on the *j*^th^ strand (template or non-template) in the *k*^th^ expression group (nascent expression strata defined above, n=6). Mutation and sequence composition counts were summed (aggregated) over *ijk* for each gene for a tumour, and then the sum of focal tumours used to calculate the observed mutation rate for each instance of *ijk*. To adjust for the expected number of upstream template strand genic lesions (expected upstream lesions), the positional category *i* for each window in each gene in each tumour was replaced with an estimated number of upstream lesions. The estimate was calculated by multiplying the trinucleotide composition of the upstream template strand genic interval by the genome-wide mutation rate for the 64 trinucleotide contexts derived from that tumour. The estimate was rounded to one decimal place and used as a replacement index *i* for the aggregation of mutation rates. For plotting and modelling only the first (5’ most) 40 spatial (0-200 kb) windows and compositionally adjusted (0-4 upstream mutations) windows were included.

### Modelling transcription coupled repair

We defined a probabilistic model of lesion detection by RNA Pol-II (variable parameter *Pd*), and its subsequent re-initiation (*Pr*) or dis-association (1-*Pr*). The model also incorporated variables for the fraction of lesions that are visible to TCR (*Pv*; mutation spectra suggest some lesions types or contexts are not subject to TCR) and a multiplier parameter (*m*) to translate experimental measurements of nascent TPM (nTPM), into number of transcripts between mutagenesis and DNA replication. Experimental measures of median nascent expression for the six expression strata were (0, 0.49, 1.16, 2.07, 3.14, 11.15 nTPM). For modelling purposes, genes had a uniform length and each had an average of 4 lesions on their template strand, allowing direct comparison to observed/expected (OE) measures of mutation rate through gene bodies (**Fig. 4g**).

With a Markov chain formulation of the lesion count dynamics, for fixed values of the independent variables (*Pd, Pr, Pv, m*), we were able to produce the expected number of mutations over 40 consecutive windows through the gene body, for each of the six expression strata. For model fitting, minimising the distance (sum root mean squared deviation, RMSD) between those 6×40 measures and the equivalent experimentally determined measures was the optimisation criterion. Parameter space was initially explored as a grid-search. Probabilities *Pd, Pr*, and *Pv* were bounded at min=0, max=1 with steps of 0.01. The expression multiplier *m* was bounded at min=0.25, max=64 with steps of 0.25. This range implies between 0.25 million and 64 million nascent Pol-II transcripts produced on average by a cell in the time between mutagenesis and DNA replication, the upper bound set generously assuming 180,000 chromatin associated RNA Pol II complexes per cell^62^, all polymerases are continuously actively transcribing, and only transcribing annotated genes, an average transcription rate of 2 kb min^-1^ in mouse liver^63^, median gene length 60 kb, and average 2,280 minutes between mutagenesis and DNA replication estimated from cell-cycle times of DEN mutagenised rat hepatocytes^64^. Grid-optimal parameters were provided as the starting point for optimisation implemented in the R optim function^65^ with default parameters to return the final optimised parameter values.

To identify the boundaries of high-confidence parameter fitting, the observed OE rates for the six expression strata were re-calculated from the bootstrap sampling of genes (sampling with replacement to original gene list size, n=1,000 replicates). The mathematical TCR model (**Supplementary File 1**) was independently optimised by grid-search for each bootstrap replicate, the empirical 95% confidence intervals reported for each model parameter.

### Mouse colony management

Animal experimentation was carried out in accordance with the Animals (Scientific Procedures) Act 1986 (United Kingdom) and with the approval of the Cancer Research UK Cambridge Institute Animal Welfare and Ethical Review Body (AWERB). Animals were maintained using standard husbandry: mice were group housed in Tecniplast GM500 IVC cages with a 12 h:12 h light:dark cycle and ad libitum access to water, food (LabDiet 5058), and environmental enrichments.

### ATAC-seq

Liver samples from P15 mice (matching the developmental time of mutagenesis) were isolated and flash frozen. ATAC-seq was performed as described previously^66^, with minor modifications to the nuclear isolation steps (in Step 1, 1 ml of 1× homogeniser buffer was used instead of 2 ml; in Step 4, douncing was performed with 30 strokes instead of 20). Pooled libraries were sequenced on a NovaSeq6000 (Illumina) to produce paired-end 50 bp reads, according to manufacturer’s instructions. Experiments were performed with 3 biological replicates.

### ATAC-seq data processing and analysis

ATAC-seq data processing was performed using a Snakemake pipeline (v.6.1.1)^67^. Adaptor sequences were removed using cutadapt (v.2.6)^68^. Reads were aligned to the reference genome (Ensembl v.91: C3H_HeJ_v1^55^) using BWA (v.0.7.17)^69^. Data from multiple lanes were merged prior to deduplication; duplicates were marked using Picard (v.2.23.8)^70^. Reads overlapping ARC regions were removed using samtools (v.1.9). Reads aligning to mitochondrial DNA were excluded from further analysis. Read positions aligning to forward and reverse strands were offset by +5 bp and -4 bp, respectively, to represent the middle of the transposition event, as described previously^71^. ATAC-seq peaks were called using MACS2 (v.2.1.2)^72^ on pooled data containing all replicates. Single nucleotide-resolution chromatin accessibility was measured and plotted as coverage of ATAC-seq ‘tags’ (Tn5 insertion sites, adjusted to represent middle of transposition event, as described above).

ATAC-seq data are available from Array Express at EMBL-EBI under accession E-MTAB-11780.

### Nucleosome positioning analysis

We used nucleosome positions determined through chemical profiling of mouse embryonic stem cells^38^ using a nucleosome centre positioning (NCP) score to signify the prevalence of nucleosome dyads for a given genomic position. We transferred genome coordinates from mm9 to mm10 using UCSC liftover^73^, before using halLiftover (v.2.1) to derive expanded C3H-specific coordinates, considering only unique non-overlapping and syntenic positions. The top 4 million dyad positions were selected based on the NCP score.

The positions and span of the major groove (either facing out or into the histones relative to the dyad) was calculated with the centre of the major groove facing inwards repeating every ±10.3 bp away from the dyad position, and spanning 5.15 bp^10^.

### CTCF ChIP-seq

Livers from P15 mice (matching the developmental time of mutagenesis) were perfused *in situ* with PBS and then dissected, minced, cross-linked using 1% formaldehyde solution for 20 min, quenched for 10 min with 250 mM glycine, washed twice with ice-cold PBS, and then stored as tissue pellets at –80°C. Tissues were homogenised using a dounce tissue grinder, washed twice with PBS, and lysed according to published protocols^74^. Chromatin was sonicated to an average fragment length of 300 bp using a Misonix tip sonicator 3000. To negate batch effects and allow multiple ChIP experiments to be performed using the same tissue, we pooled ten livers for each experiment; 0.5 g of washed homogenised tissue was used for each chromatin immunoprecipitation, using 20 μg CTCF antibody (rabbit polyclonal, Merck Millipore 07-729, lot 2517762). Library preparation was performed using immunoprecipitated DNA or input DNA (max 50 ng) as described previously^75^ with the ThruPLEX DNA-Seq library preparation protocol (Rubicon Genomics, UK). Libraries were quantified by qPCR (Kapa Biosystems), and fragment size was determined using a 2100 Bioanalyzer (Agilent). Pooled libraries were initially sequenced on a MiSeq (Illumina) to ensure balanced pooling, followed by deeper sequencing on a HiSeq4000 (Illumina) to produce paired-end 150 bp reads, according to manufacturer’s instructions; only HiSeq libraries were used for downstream analyses. Experiments were performed with five biological replicates.

To identify ChIP-seq positive regions, we trimmed the HiSeq sequencing reads to 50 bp and then aligned them using BWA (v.0.7.17) using default parameters. Uniquely mapping reads were selected for further analysis. Peaks were identified for each ChIP library and input control using MACS2 (v.2.1.2) callpeak with default parameters, and all peaks with a q-value >0.05 were included in downstream analyses. Input libraries were used to filter spurious peaks associated with a high input signal using the GreyListChIP R package^76^. Biologically-reproducible peaks were identified by merging ChIP-seq peaks defined as above from individual replicates and selecting those that overlapped ≥2 individual replicate peaks.

ChIP-seq data are available from Array Express at EMBL-EBI (accession ID pending).

### Transcription factor binding site identification and analysis

ChIP-seq data for transcription factors, apart from CTCF (see above), were obtained from Life Science Database Archive (https://dbarchive.biosciencedbc.jp/datameta-list-e.html) with genomic coordinates for the mm9 reference assembly. Liver-specific ChIP-seq was used whenever possible, otherwise files marked with “All cell types’’ were used instead (**Supplementary Table 2**). Genomic coordinates were lifted to mm10 using liftOver, and then lifted to the C3H genome assembly using halLiftover (as above). Overlapping ChIP-seq regions were merged, using the outermost coordinates as the new start/end of regions. FASTA sequences of the regions were extracted using Bedtools getfasta (v2.27.1) and used together with non-redundant vertebrate position weight matrices from JASPAR^77^ to run FIMO (MEME suite)^78^ with default parameters to detect motifs within ChIP-seq peaks. Those motifs were then filtered based on an overlap with ATAC-seq peaks (defined above) to ensure that the analysed set was within open chromatin regions of P15 C3H mouse livers. For CTCF binding site analysis, in-house generated ChIP-seq data (described above) was used. For wider flank (1 kb) analysis, all motifs (JASPAR matrix profile MA0139.1) within the peaks were retained regardless of ATAC-seq intersection, allowing multiple motifs per ChIP-seq peak.

For high-resolution CTCF and TFBS analysis (**Fig. 6**) only one highest-scoring motif per ChIP-seq peak was retained. Similarly, for aggregate transcription factor analysis, only one highest-scoring motif per ChIP-seq peak was retained if it overlapped with ATAC-seq peak. Total of 129 transcription factors were analysed based on ChIP-seq and position weight matrix (PWM) availability, RNA-seq support for transcription factor expression (≥1 TPM) in P15 mouse liver^2^. In all the analyses ‘bit score’ refers to the information content of the whole position. Within the motif, only mutations with the reference nucleotide matching consensus nucleotide from PWM were retained. In the flanks mutations from all reference nucleotides were used.

### CTCF structural analysis

High resolution crystal structures for CTCF zinc fingers complexed with binding site DNA were obtained from the Protein Data Bank (PDB:5YEL, PDB:5T0U, PDB:5UND)^79,80^. As no single structure contains all 11 CTCF zinc fingers, a composite structure was compiled through alignment using PyMOL (v.2.5.2)^81^ align function. PDB:5UND A chain 406-556 was aligned to PDB:5T0U A chain (root mean square deviation: 1.06Å); then PDB:5YEL A chain aligned to PDB:5UD chain A (root mean square deviation: 1.3Å). The composite image (**Fig. 6a**) then shows PDB:5T0U A chain 289-405, PDB:5UND A chain 406-488, and PDB:5YEL A chain 489-556 which collectively spans CTCF zinc fingers 2 to 11 inclusive. The bound DNA strands comprise PDB:5YEL F chain 1-24, PDB:5T0U C chain 7-23, PDB:5T0U B chain 1-18, PDB:5YEL E chain 5-26.

Protein:DNA contact distance measurements were performed using the Protein Contacts Atlas^82^. Non-covalent interatomic contacts of ≤3Å between CTCF protein and DNA were considered close contacts. Close contacts of atoms within phosphate groups or deoxyribose were considered backbone, other DNA contacts annotated as base contacts. Close base contacts involving atoms expected to acquire DEN induced mutagenic adducts^20^ or structurally equivalent positions in other bases (purines: N6, O6; pyrimidines: O4, N4, O2) were annotated as lesion site contacts. Distance measurements were taken separately for each structure (rather than from the composite) and excluded PDB:5T0U nucleotide contacts upstream of binding motif position +1 where this structure substantially deviates from PDB:5YEL. PDB:5T0U is truncated at zinc finger 7 whereas PDB:5YEL extends to zinc finger 11 and makes additional base-specific contacts absent from PDB:5T0U. Close backbone, base, and lesion site contacts were reported if the distance threshold criteria were met in any of the 3 considered structures, though concordance was high in the overlapping regions.

### Histology and image analysis

Digitised histology images of DEN-induced tumours^2^ were obtained from Biostudies (accession S-BSST383): https://www.ebi.ac.uk/biostudies/studies/S-BSST383.

Whole slide images (WSIs) of tumours that met inclusion criteria (cellularity >50% and DEN1 signature >80%) were annotated in QuPath (v.0.2.2)^83^ using the polygon tool to include neoplastic tissue and exclude adjacent parenchyma, cyst cavities, processing artefacts, and white space. For tumours with multiple transections, only a single WSI was used.

Annotations were reviewed for quality by two Histopathologists (J.C. and S.J.A.). Using Groovy in QuPath, annotated regions were tessellated into fixed size, non-overlapping 256 µm^2^ tiles. For segmentation of epithelioid nuclei, a pre-trained StarDist^84^ model (he_heavy_augment.zip) was downloaded from https://github.com/stardist/stardist-imagej/tree/master/src/main/resources/models/2D and an inference instance was deployed using Groovy across the tiles in QuPath, built from source with Tensorflow^85^, with a minimum detection threshold of 0.5. Python (v.3.9.7) was used for downstream analyses. Data were filtered to exclude extreme outliers: Tiles with ≤43 nuclei per tile were excluded; nuclei with area ≥227.18386 µm, circularity of ≤0.4841, or non-computable circularity were excluded. From the 245 WSI (n=237 mutationally asymmetric tumours and n=8 symmetric tumours), 70,414 tiles were generated, and 9,999,783 nuclei were segmented (post-filtering). To compute inter-nuclear distance, for each nucleus in a tile represented by its X-Y centroid coordinates, nearest neighbours were identified using the k-dimensional tree function from the spatial module of SciPy (v.1.7.1)^86^. The Euclidean distance for each nearest neighbour pair was computed using the paired distances function from the metrics module of SciKit-Learn (v.1.0.2)^87^. The median nuclear area, median nuclei per tile, and median inter-nuclear distances were compared between asymmetric and symmetric tumours using a two tailed Wilcoxon rank sum test.

### Symmetric versus asymmetric tumour comparison

Mutationally symmetric tumours (defined above; >99% of autosomal mutations in genomic segments with *abs(S)* < 0.2) were filtered to the subset that met the same inclusion criteria as the other n=237 tumours analysed in this study (>50% cellularity (after adjusting for the presence of two genomes) and >80% substitution mutations attributed to DEN1 signature). Eight tumours met this criteria. We subsequently show that these tumours are not whole genome duplicated, but that they contain both daughter lineages of an originally mutagenised cell (**Extended Data Fig. 5b**). For each autosomal variant in a tumour we calculated its VAF quantile position amongst point mutations in that tumour, using the R ecdf function^65^. The quantile positions (range 0-1) were grouped into consecutive bins of 0.005 unit span, i.e. the 0.995-1.0 was the right-most bin representing the top 0.5% of VAF values for mutations in a tumour. The mutations within a VAF quantile bin were classified as either overlapping or not overlapping the genomic span of the most highly expressed genes (stratum 6) using the R data.table foverlaps function^88^. The counts of overlapping and non-overlapping mutations from the focal tumour were compared as a two-tailed Fisher’s exact test to the equivalent counts aggregated from all asymmetric tumours (excluding the focal tumour in the case of asymmetric focal tumours for the calculation of background expectation). The same analysis was performed in aggregate for all symmetric tumours (n=8) compared to all asymmetric tumours (n=237). The calculations were repeated for each of the 200 consecutive bins to demonstrate the VAF range over which high VAF mutations are preferentially enriched in highly expressed genes specifically in symmetric tumours, as predicted under NER-TRIM.

### Computational analysis environment

Except where otherwise noted, analysis was performed in Conda environments and choreographed with Snakemake ^67^ running in an LSF 965 or Univa Grid Engine batch control system (**Supplementary Table 3**). Statistical tests were performed in R (v.4.0.5) using fisher.test, ks.test, cor.test, and wilcox.test functions for Fisher’s exact, Kolmogorov-Smirnov, Pearson’s and Spearman’s correlation, and Wilcoxon tests, respectively. Graphics were generated using R.

## Code and data availability

The analysis pipeline including Conda and Snakemake configuration files can be obtained without restriction from the repository https://git.ecdf.ed.ac.uk/taylor-lab/lce-si

Raw data files are available from Array Express at EMBL-EBI. ATAC-seq accession number E-MTAB-11780; ChIP-seq accession number pending.

## Key resources

The key reagents and resources required to replicate our study are listed in Supplementary Table 3. For externally sourced data, where applicable, URLs that we used can be found in the Git repository https://git.ecdf.ed.ac.uk/taylor-lab/lce-si

## Acknowledgements and funding

We thank P. Bankhead for supervision of image processing; T. Deegan for informative discussions; N. Hastie, C. Ponting, and W. Bickmore for comments on the manuscript; M. Roller for assistance with data curation; the CRUK Cambridge Institute Core facilities for their valuable contribution: CRUK Biological Resources (A. Mowbray), Genomics (P. Coupland), and Bioinformatics (G. Brown and M. Eldridge); Edinburgh Genomics, The University of Edinburgh for provision of sequencing services; and the European Molecular Biology Laboratory for access to computational resources.

This work was supported by: the MRC Human Genetics Unit core funding programme grants (MC_UU_00007/11 and MC_UU_00007/16), MRC Toxicology Unit core funding (RG94521), Cancer Research UK Cambridge Institute core funding (20412), European Molecular Biology Laboratory, ERDF/Spanish Ministry of Science, Innovation and Universities-Spanish State Research Agency/DamReMap Project (RTI2018-094095-B-I00), and the Wellcome Trust (WT202878/B/16/Z). Edinburgh Genomics is partly supported through core grants from NERC (R8/H10/56), MRC (MR/K001744/1) and BBSRC (BB/J004243/1).

M.D.N is a cross-disciplinary post-doctoral fellow supported by funding from CRUK Brain Tumour Centre of Excellence Award (C157/A27589). J.C. is supported by a Wellcome Trust PhD Training Fellowship for Clinicians (WT223088/Z/21/Z) as part of the Edinburgh Clinical Academic Track (ECAT) programme. O.P. was funded by a BIST PhD fellowship supported by the Secretariat for Universities and Research of the Ministry of Business and Knowledge of the Government of Catalonia and the Barcelona Institute of Science and Technology. V.S. was supported by an EMBL Interdisciplinary Postdoc (EIPOD) fellowship under Marie Skłodowska Curie actions COFUND (664726). S.J.A. received a Wellcome Trust PhD Training Fellowship for Clinicians (WT106563/Z/14/Z) and National Institute for Health and Care Research (NIHR) Clinical Lectureship.

For the purpose of open access, the authors have applied a Creative Commons Attribution (CC BY) licence to any Author Accepted Manuscript version arising from this submission.

## Author contributions

M.S.T. conceived the project and designed the analyses. F.C. and S.J.A. performed mouse experiments and collected tissue samples. S.J.A. performed ChIP-seq experiments. S.C. performed ATAC-seq experiments. C.J.A., L.T., J.L., M.D.N., and M.S.T. designed and performed computational analysis of genomic data; O.P., V.S. and P.A.G. provided supporting genomic analyses. J.C. and S.J.A. analysed and annotated histology images. J.C. performed computational image analysis. N.L.B., P.F., C.A.S., D.T.O., S.J.A., and M.S.T. lead the Liver Cancer Evolution Consortium and supervised the work. Research support to P.F., D.T.O., S.J.A., and M.S.T. funded the work. D.T.O., S.J.A., and M.S.T. wrote the manuscript, with contributions from C.J.A., L.T., J.L., M.D.N., and J.C.. All authors had the opportunity to edit the manuscript. All authors approved the final manuscript.

## Extended data

**Extended Data Fig.1.**
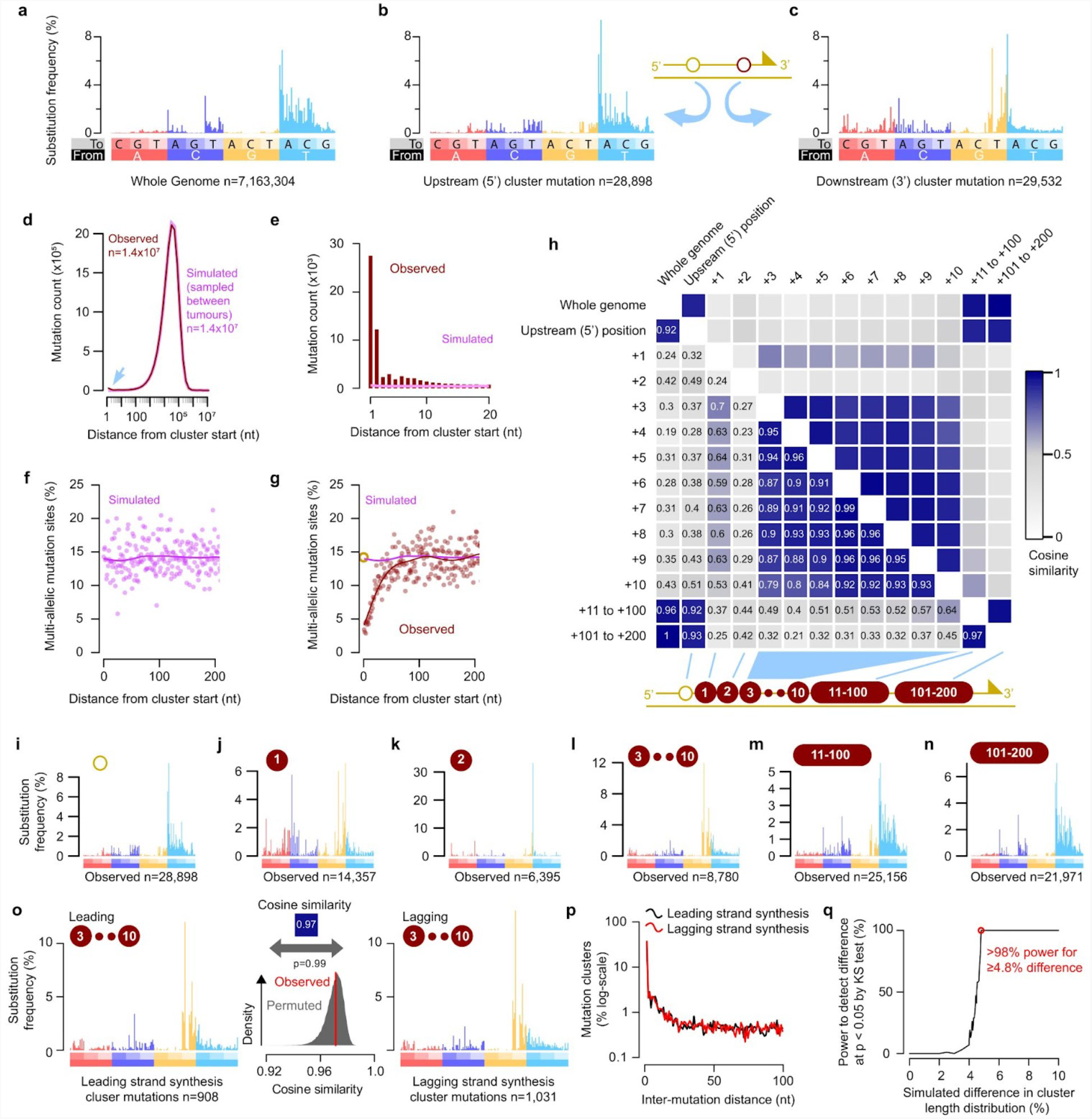
Tracts of low-fidelity replication downstream of lesion induced mutations. **a**, Genome-wide mutation signature of DEN induced tumours. **b**, Signature of mutation cluster upstream (5’) position mutations, oriented so the lesion containing strand is the replication template. **c**, Signature of downstream mutations in the cluster (2.2% of clusters have two downstream mutations). **d**, Frequency distribution of the spacing between adjacent observed (dark-red) and simulated (pink) mutations for all tumours (n=237). The simulated data were generated by sampling mutations across all other tumours to create proxy tumour datasets with identical mutation counts. Excess clustering of observed mutations (blue arrow) accounts for only 0.4% of the total mutation burden; observed and simulated distributions otherwise overlap. **e**, Frequency distribution for spacing between mutations, corresponding to the closely spaced mutations highlighted (blue arrow) in **d. f**, Multiallelism is a hallmark of lesion templated mutations^2^. The multiallelic rate (y-axis, fraction of mutation sites with multiallelic variation) for simulated data (pink spots). Curve shows best-fit spline (25 degrees of freedom) for the downstream mutations. **g**, As for **f** but showing observed data (red), demonstrating a pronounced and specific depletion of multiallelic variation immediately downstream of the cluster start (yellow circle). **h**, Heatmap summarisising cosine similarity between mutation clusters with different inter-mutation spacing (schematic in lower panel). Upstream cluster mutations closely match the genome wide mutation spectrum. Mutations 3 to 10 nt downstream of the 5’ mutation share a common signature. **i-n**, Mutation signature profiles for clustered mutations; distance from the upstream mutation (number in brown circle) relate to schematic in **h**. Mutation counts in each category indicated below the plot. **o**, The mutation spectrum of downstream mutations closely matches between leading and lagging strand replication. The observed cosine similarity between mutation spectra is robustly within the range expected by random permutation of mutations between leading and lagging strands (n=10^5^ permutations, two tailed empirical p=0.99). **p**, The distribution of mutation cluster length also matches between leading (black) and lagging (red) strands. **q**, Simulations show >98% power to detect a ≥4.8% difference in the distribution of cluster lengths.

**Extended Data Fig.2.**
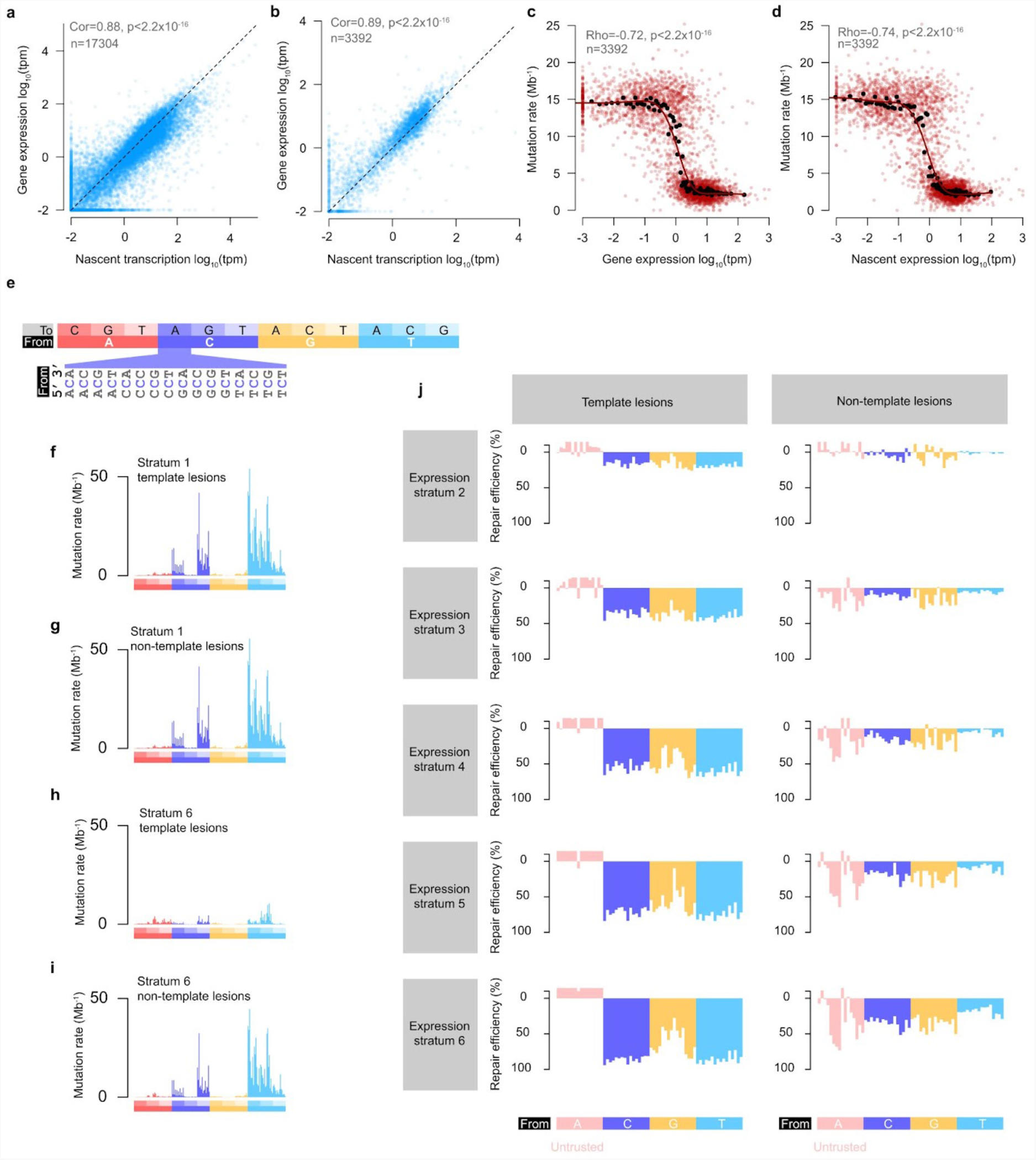
Transcription and lesion repair have strand-specific, expression-dependent mutation signatures. **a**, Mature transcript expression and nascent transcription (intron mapping RNA-seq reads) are highly correlated; one point per gene. **b**, As for panel **a** but restricted to the genes spanning in aggregate across tumours >2 million nucleotides of strand resolved tumour genome (n=3,392). **c**, Mature transcript gene expression (x-axis) negatively correlates with composition normalised mutation rate (y-axis) where lesions are on the transcription template strand (one red point per gene). Red curve shows the best-fit spline (8 degrees of freedom) through the red points. Black points show gene expression measures for centile bins of gene expression. **d**, As for **c**, but x-axis shows nascent RNA estimates of transcription. P-values for panels **a-d** are too small to precisely calculate (p<2.2×10^−16^). **e**, Nucleotide order used for 192 category mutation spectra in panels **f-i**. Expanded segment shows the flanking nucleotide context for C→A mutations; the same ordering of flanking nucleotides is used for all mutation types. **f-i**, Mutation rate spectra for non-expressed (stratum 1) genes are closely matched for template (**f**) and non-template (**g**) lesion strands. For highly expressed genes (stratum 6), the mutation rate is reduced for both strands and the spectrum differs between template strand (**h**) and non-template strand (**i**) lesions. **j**, The profile of lesion repair efficiency differs between template strand lesions and non-template strand lesions of expressed genes. Repair efficiency is calculated as the percent change in mutation rate for a trinucleotide sequence context (n=64 categories) relative to the average for both strands in non-expressed genes (stratum 1). The y-axis is inverted to indicate reduction in mutation rate from increased repair. Transcription coupled repair shows similar efficiency for C and T lesions on the template strand. Transcription associated repair on the non-template strand shows preferential repair of C lesions compared to T lesions. Mutations from apparent A lesions (and to a lesser extent G lesions) are rare and, as shown in subsequent sections, should not be evaluated as lesions on the indicated nucleotide, but are included for completeness (y-axis values < -10 truncated).

**Extended Data Fig.3.**
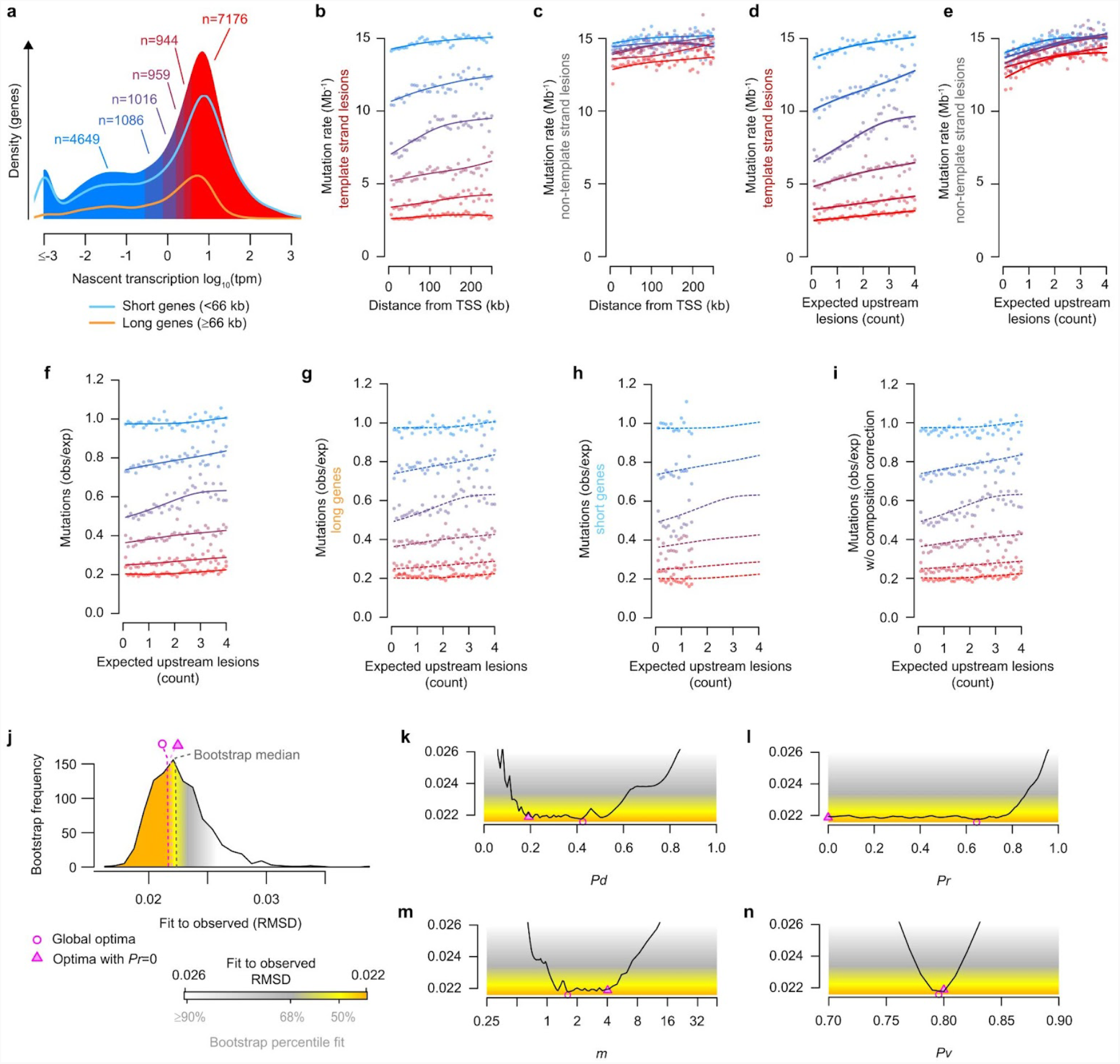
Transcription coupled repair through gene bodies. **a**, Distribution of genes by measure of nascent gene expression. Blue→red denote increasing categories of nascent gene expression (thresholds as per **Fig. 3b**). Density shown for all genes included in gene-body analysis (n=15,830) and scaled density curves shown for the long-gene (≥66 kb, orange) and short-gene (<66 kb, blue) component subsets. **b**, Mutation rates for genes with template strand lesions. Genes classified by expression strata and mutation rates calculated in 5kb consecutive windows from the TSS. Curves show best-fit splines (3 degrees of freedom). **c**, As for **b** but considering genes with non-template lesions. **d**, Analysis of panel **b** but distance from TSS converted to expected upstream lesion count (x-axis) by per-tumour normalisation (**Fig. 4b**). **e**, as for **d** but showing genes with non-template lesions (data from **c**). **f**-**i**, Observed versus expected mutations for each expression strata, calculated as the ratio of template strand (panel **d**) to non-template (panel **e**) strand mutations, after adjusting for expected upstream lesions. Lower values indicate greater transcription coupled repair. All genes (**f**; also shown in **Fig. 4c**), long-genes only (**g**), short-genes only (**h**); all genes without applying mutation rate corrections for sequence composition (**i**); dashed curves show the best-fit splines from panel **f** for comparison. **j**, Frequency distribution of the distance (root mean square deviation, RMSD) between the observed profile of transcription coupled repair efficiency (panel **f**) and bootstrap samples (n=1,000) of that data, to serve as a comparator for model fitting to the observed data. Shading denotes a distance ≥90% bootstraps (white) to 68% (grey), to 50% (yellow) and grid-search optimum (orange). **k-n**, Summary of TCR model fitting grid-search parameter space. For each tested value of model parameters *Pd, Pr, m* and *Pv* the distance (RMSD) of optimal fit is plotted (black curve). Shading defined in **j**. Optimal solutions identified by gradient descent indicated by pink shapes (circle *Pr*≥0, triangle *Pr*=0).

**Extended Data Fig.4.**
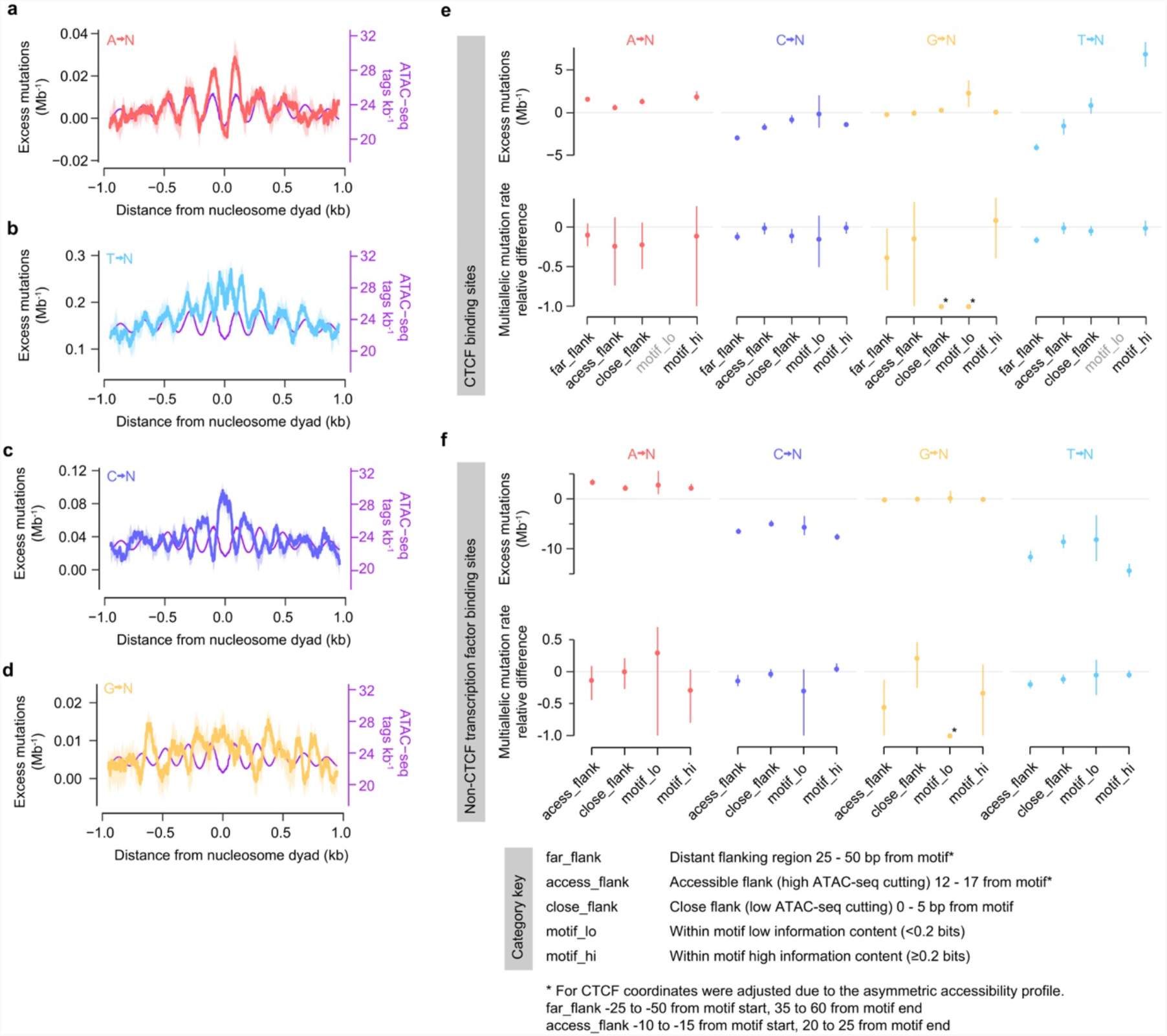
Lesion induced mutation patterns at DNA:protein interaction sites. **a**, Excess mutations resulting from A lesions in accessible DNA (relative to the genome-wide trinucleotide mutation rate) centred on the nucleosome dyad. DNA accessibility as measured by ATAC-seq (purple, higher values mean more accessible chromatin). **b-d**, Relative mutation rates as **a**, for apparent T lesions (**b**), C lesions (**c)**, and G lesions (**d**); in each case, except A→N mutations, the mutation rate is lower in accessible DNA and higher in less-accessible DNA. **e**, Mutation rates and multiallelic rates for sequence categories (Methods) within, and adjacent to, CTCF binding sites, stratified by the identity of the inferred lesion containing nucleotide. Point estimate (circles) and bootstrap 95% confidence intervals (whiskers) are shown for the rate difference relative to genome-wide expectation (y=0, mutations Mb^-1^ for mutation rates, relative difference metric for multiallelic variation). All rates are adjusted for trinucleotide composition. Instances where the motif_lo category has too few observed or expected mutations to calculate estimates (x-axis label grey) have no data point. Where the observed level of multiallelic variation is zero (asterisk) bootstrap confidence intervals cannot be calculated. **f**, Mutation rates and multiallelic variation for P15 liver expressed transcription factors; plots as in (**e**).

**Extended Data Fig.5.**
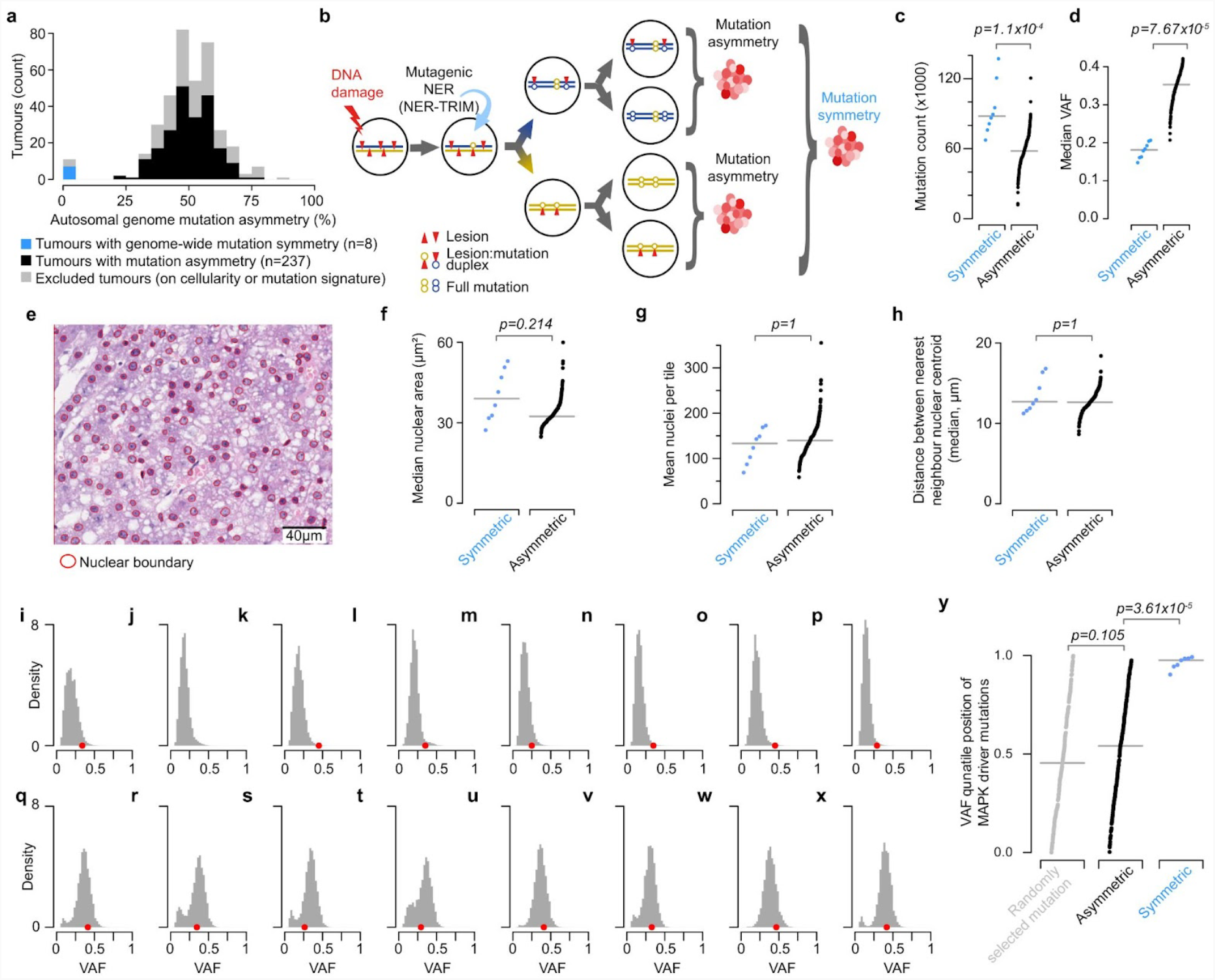
Mutagenic nucleotide excision repair. **a**, Most DEN induced tumours show pronounced mutation asymmetry across approximately 50% of their genome. Asymmetric tumours meeting inclusion criteria (mutation signature and cellularity thresholds; black) are included in the preceding analyses of this study. In addition, here we include a subset of tumours that were excluded due to the absence of mutation asymmetry (n=8, blue). **b**, The mutational symmetry of these tumours could be explained if both daughters of the originally mutagenised cell persist (schematic). Mutagenic NER in the first generation of the mutagenised cell could produce mutations at the same base pair in both daughter lineages; such mutations would have approximately double the variant allele frequency (VAF) of mutations confined to one daughter lineage. Whole genome duplication in the first generation of the mutagenised cell could also produce symmetric tumours (schematic not shown). **c**, Tumours with symmetric mutation patterns have a significantly higher mutation load than those with asymmetric mutations, consistent with mutations from both mutagenised strands contributing to the tumour. Statistical analysis (p=1.1×10^−4^) by two tailed Wilcoxon rank sum test. In panels **c**,**d**,**f**,**g**,**h** points are individual tumours, bar is median, statistical tests are based on n=8 symmetric and n=237 asymmetric tumours, all reported p-values are Bonferroni corrected (n=5 tests). **d**, The median VAF for mutations in symmetric tumours is approximately half that of asymmetric tumours. Statistical analysis (p=7.67×10^−6^) by two tailed Wilcoxon rank sum test. **e**, Automated nuclear detection (red circles) and quantification in an exemplar hematoxylin and eosin stained tumour section (93131_N2). Original digitised magnification x200; scale bar indicated. **f**, Nuclear area is not significantly different between symmetric and asymmetric tumours (p=0.215, two tailed Wilcoxon rank sum test), indicating similar DNA content and arguing against mononuclear whole-genome duplication. **g**, The density of nuclei is not significantly different between symmetric and asymmetric tumours (p=1, two tailed Wilcoxon rank sum test), arguing against both mononuclear and possibly multi-nuclear whole genome duplication. **h**, Internuclear distance is not significantly different between symmetric and asymmetric tumours (p=1, two tailed Wilcoxon rank sum test), arguing against multi-nuclear whole genome duplication. **i-p**, VAF frequency distributions for symmetric tumours, indicating the VAF of MAPK pathway driver mutations (red points, also in **q-x**). For symmetric tumours, the driver VAFs are strongly right-biassed. This is consistent with mutagenic NER copying the same driver mutation site into both daughter genomes of the mutagenised cell, and in turn both daughter lineages (containing either the same driver mutation, or multiallelic driver mutations at the same site) contributing to the resultant tumour. **q-x**, VAF frequency distributions for example asymmetric tumours. **y**, MAPK pathway driver mutations are biassed to the highest VAF values in symmetric tumours but not in asymmetric tumours (p=3.61×10^−5^ two tailed Wilcoxon rank sum test, Bonferroni corrected). VAF quantile position (y-axis) indicates the fraction of mutations in a tumour that have lower VAF than the driver mutation (quantile of 1.0 indicates all other mutations in that tumour have a lower VAF). Horizontal bars indicate median VAF quantile position of the focal driver mutations. As a null expectation for comparison, one mutation was randomly selected from each of the asymmetric tumours (grey points).

**Supplementary Table 1** | **Table of tumours sequenced containing key metadata** (Excel file).

**Supplementary File 1** | **Mathematical model for transcription coupled repair** (PDF).

**Supplementary Table 2** | **Table of ChIP-seq transcription factors and tissues of origin from ChIP-Atlas database** (Excel file).

**Supplementary Table 3** | **Table of key resources and software** (Excel file).

