## Supplementary Information for "Strand-resolved mutagenicity of DNA damage and repair"

Supplementary File 1: Mathematical model for transcription coupled repair

#### Document overview

This document provides the mathematical details for the analytic model of transcription coupled repairs - 'McTcr'. R code for the model can be found in the file tcrMmMn.R; relevant parts of this code - linking the mathematical expressions to the code - are given below.

#### Method overview

Our aim is to establish a mathematical model, linking fundamental parameters of TCR, to the aggregate mutational pattern seen over genes and individual tumours.

As for a given gene the mutation pattern is influenced by genic sequence composition and individual tumour mutational burden, we compared observed:expected mutation rates normalised to the expected number of upstream lesions, see Fig 4 in the main text. Using the 'gene distance' variable of expected number of upstream lesions, all genes can be treated identically - except for gene length (in units of expected upstream lesions) and expression (modelled as the number of RNApols that initiate transcription prior to the first cell division of the original mutagenised cell). Hence, we consider how the mutational profile for a given gene alters as transcription coupled repair proceeds; the aggregrate profile expected to be seen over all tumours, for all genes of a given length and having seen $n$ RNApols initiate transcription, may then be acquired for a given set of model parameters by summing over the single gene mutational profie.

#### Model and methods strategy

We take a gene body of length $l$ which may have lesions placed upon it. We assume the number of lesions up to a position $x$ in the gene is distributed as $\text{Pois}(\mu x)$, where $\mu$ is the initial lesion rate; we keep this in the model for generality however as we measure gene position in units of 'expected upstream mutations', $\mu = 1$ when we compare to the data. We assume $n$ RNA polymerases begin transcription at the TSS. These polymerases are ordered (first RNApol starts transcription then second etc). If a polymerase encounters a lesion, the lesion is detected with probability $p_d$. If detected, it is repaired. After repair the polymerase may restart with probability $p_r$, if not it falls from the strand. We also assume that a certain proportion of lesions are invisible to the RNA polymerases; each lesion is visible with probability $p_v$. We are interested in the number of remaining lesions after $n$ RNA polymerases have exited the gene. See Fig 4 in the main text for a graphically summary of the mode.

Our aim is to determine the expected number of lesions that remain after $n$ RNA polymerases have exited the gene, in any increment of the gene body (e.g. $[x_1, x_2] \subset [0, l]$). To do so, we pick a position $x$ in the gene and then investigate the probability we have $j$ lesions remaining after $n$ RNA polymerases have exited the gene given that we start with $i$ lesions before $x$. This amounts to deriving the transition probabilities in a specific Markov chain. From the transition probabilities, we can obtain the expected number of remaining lesions after $n$ polymerases, given we start with $i$ initial lesions. As the number of initial lesions before $x$ is a Poisson random variable, we then average over the initial lesion number to obtain the mean remaining lesions before $x$. From the mean we can deduce the expected number of lesions remaining in any increment in the gene, and over multiple genes. The expected number of lesions with repair, compared to the expected number of lesions in the absence of any repair is comparable to the experimental data, and is given by Equation (8) below. The derivation of Equation (8), and how to numerically evaluate the expression is the purpose of this document.

#### Probability transition matrix

We take a position $x$ in the gene, and suppose we have $i$ initial lesions before $x$. For now assume all lesions are visible, that is $p_v = 1$. We now sequentially send through polymerases and let $L_n$ denote the number of remaining lesions after $n$ polymerases have exited the gene, so here $L_0 = i$. We seek $\Pr(L_n = j|L_0 = i)$. As $L_n$ is a Markov chain, it is sufficient to consider $\Pr(L_1 = j|L_i = i)$, from which $\Pr(L_n = j|L_0 = i)$ can be obtained by appropriate matrix multiplication according to standard Markov chain theory. We turn to $\Pr(L_1 = j|L_0 = i)$.

The strategy is to first mark each of the $i$ initial lesions depending on whether the polymerase will detect these, i.e. we binomially thin the lesions, and then consider how many of the thinned lesions the polymerase will detect before exiting the increment of interest (falling of the gene or passing beyond the position $x$).

Let $L'_0$ be the number of lesions the RNApol could detect before $x$, so $L'_0$ is binomial with $i$ trials and with success probability $p_d$. Then the number of lesions repaired by the first polymerase, $R_1$ is geometric with parameter $q_r = 1 - p_r$ truncated to be within $1, \ldots, L'_0$. So for $k = 1, \ldots, L'_0$

$$\Pr(R_1 = k) = \begin{cases} p_r^{k-1} q_r, & k < L'_0 \\ p_r^{L'_0 - 1}, & k = L'_0 \end{cases}$$

If $L'_0 = 0$, then $\Pr(R_1 = 0) = 1$.

If $L_1 = j$, then $i - j$ lesions have been repaired, so

$$\Pr(L_1 = j|L_0 = i) = \Pr(R_1 = i - j) = \sum_{y=0}^{i} \Pr(R_1 = i - j|L'_0 = y)\Pr(L'_0 = y). \tag{1}$$

We introduce the indicator $1(A)$ which returns 1 when the condition $A$ is true and 0 otherwise. Notice that no lesions are repaired only if all lesions are undetected, that is $\Pr(L_1 = i|L_0 = i) = \Pr(L'_0 = 0)$. Hence separating out this case, and assuming $j \leq i$,

$$\begin{aligned} \Pr(L_1 = j|L_0 = i) &= 1(j = i)\Pr(L'_0 = 0) + 1(j < i)\sum_{y=1}^{i} \Pr(R_1 = i - j|L'_0 = y)\Pr(L'_0 = y) \\ &= 1(j = i)\Pr(L'_0 = 0) + 1(j < i)\sum_{y=1}^{i} (p_r^{i-j-1} q_r 1(i - j < y) \\ &\quad + p_r^{y-1} 1(i - j = y)) \binom{i}{y} p_d^y q_d^{i-y}. \end{aligned}$$

We can extend this to $\Pr(L_1 = j|L_0 = i)$ for $j > i$ by setting the probability to 0 in this case.

Let $P$ be the matrix with elements $P_{i,j} = \Pr(L_1 = j|L_0 = i)$, and $P^{(k)}$ be $P$ multiplied by itself $k$ times. Then from standard Markov chain theory we identify

$$\Pr(L_n = j|L_0 = i) = P_{i,j}^{(n)}.$$

If $L_0$ is known then $P$ would be of size $L_0 \times L_0$ as the number of lesions before $x$ can take values in $\{0, \ldots, L_0\}$. However, as we assume $L_0$ is Poisson, then $L_0$ is in principle unbounded. Momentarily, to be justified shortly below, assume $L_0 \leq L_{\max}$ for some integer $L_{\max}$. Then $P$ would be of size $L_{\max} \times L_{\max}$.

R code to construct this one step transition probability matrix $P$ for fixed $p_d$, $p_r$ is as follows:

```r
#function to populate elements of one step transition matrix
trans_elements = function(i,j,pd,pr){
  qr = 1-pr

  truei = i-1
  truej = j-1

  if (truei<truej){
    toreturn = 0

  } else if (truei==truej){

    toreturn = dbinom(0,truei,pd)

  } else if (truei>truej){

    summands = sapply(1:truei,function(y){
      if (truei - truej <y){
        relsummand = (pr^(truei-truej-1)*qr)*dbinom(y,truei,pd)
      } else if (truei-truej == y){
        relsummand = pr^(y-1)*dbinom(y,truei,pd)
      } else if (truei - truej > y){
        relsummand = 0
      }
      return(relsummand)
    })

    toreturn = sum(summands)
  }

  return(toreturn)
}


#matrix (i+1,j+1)th element
#probability have j remaining lesions after poly exits given started with i lesions
get_onestep_transmatrix = function(Lmax,pd,pr){

  Lmaxp1 = Lmax+1
  qd = 1-pd
  qr = 1-pr

  #populate 1 step transition matrix
  trans_matrix = matrix(0,nrow = Lmaxp1,ncol = Lmaxp1)
  for (i in 1:Lmaxp1){
    for (j in 1:Lmaxp1){
      trans_matrix[i,j] = trans_elements(i,j,pd,pr)
    }
  }
  return(trans_matrix)
}
```

#### Expected number of lesions remaining

We again concern ourselves with the lesions remaining before $x$. From the above, with fixed $L_0$, we can obtain $\Pr(L_n = j|L_0)$, and thus determine the expected number of lesions remaining

$$\mathbb{E}[L_n|L_0] = \sum_{j=0}^{L_0} \Pr(L_n = j|L_0) j.$$

Our primary aim is

$$\mathbb{E}[L_n] = \sum_{i=0}^{\infty} \mathbb{E}[L_n|L_0 = i] \Pr(L_0 = i) \tag{2}$$

with $L_0$ distributed as a Poisson variable with mean $\mu x$. Equation (2) contains an infinite sum, however for some precision parameter $\delta > 0$, we may choose $L^\delta_{\text{max}}$ such that

$$\mathbb{E}[L_n] - \sum_{i=0}^{L^\delta_{\text{max}}} \mathbb{E}[L_n|L_0 = i] \Pr(L_0 = i) < \delta,$$

due to the following argument.

First notice that the number of lesions is monotone decreasing in $n$. Hence for any $n$, $\mathbb{E}[L_n|L_0 = i] \leq i$, and so,

$$\mathbb{E}[L_n] - \mathbb{E}[L_n 1(L_n \leq L^\delta_{\text{max}})] = \sum_{i=L^\delta_{\text{max}}+1}^{\infty} \mathbb{E}[L_n|L_0 = i) \Pr(L_0 = i) \tag{3}$$

$$\leq \sum_{i=L^\delta_{\text{max}}+1}^{\infty} i \Pr(L_0 = i) \tag{4}$$

$$= \mathbb{E}[L_0] - \sum_{i=0}^{L^\delta_{\text{max}}} i \Pr(L_0 = i). \tag{5}$$

Thus if we wish to compute $\mathbb{E}[L_n]$ with an error of at most $\delta > 0$, we choose $L^\delta_{\text{max}}$ such that

$$\mu x - \sum_{i=0}^{L^\delta_{\text{max}}} i \Pr(L_0 = i) \leq \delta,$$

which may be found numerically. If we wish to evaluate $\mathbb{E}[L_n]$ at a grid of positions $x_1$, $x_2 \ldots, x_y$, then we can carry out the procedure above for each $x_i$, obtaining a sequence of $L^\delta_{\text{max}}$ and then select the maximum of these. When comparing with the experimental data, we selected $\delta = 10^{-5}$, and work with

$$\sum_{i=0}^{L^{\delta=10^{-5}}_{\text{max}}} \mathbb{E}[L_n|L_0 = i] \Pr(L_0 = i).$$

Until now we have assumed all lesions are visible. However if $p_v < 1$, then by Poisson thinning the arguments presented in above holds identically but with

$$\mathbb{E}[L_n] \mapsto \mu x(1 - p_v) + \mathbb{E}[L_n],$$

and $L_0 \sim \text{Pois}(\mu x p_v)$.

To evaluate $\mathbb{E}[L_n]$ numerically we use the following R code. Note that we wish to evaluate $\mathbb{E}[L_n]$ at a number of $x$ positions, this is the variable 'upsteam grid'. Further we set $\mu = 1$ as we work in units of expected upstream mutations.

```
library(magrittr)
#get the cumulative number of lesions before the positions
#in upstream grid.
get_cmllesions_MC = function(pd,
                             pr,
                             npoly,
                             upstream_grid,
                             Lmax,
                             pv){




  trans_matrix = get_onestep_transmatrix(Lmax,pd,pr)
  #now multiple one step transition matrix to get k step transition matrix
  list_iterated_transmatrix = list()
  list_iterated_transmatrix[[1]] = trans_matrix
  for (k in 2:npoly){
    list_iterated_transmatrix[[k]] = list_iterated_transmatrix[[k-1]] %*% trans_matrix
  }

  #get expected values for different initial conditions
  #the elements of this list are matrices
  #matrix k is if we start with k lesions

  mat_ep_lesionnum_detinitial = lapply(1:npoly, function(k) list_iterated_transmatrix[[k]][1:(Lmax+1),1

#now average over the initial number of lesions with truncated Poisson
  mat_epcml_lesionnum =  lapply(1:npoly, function(x){

    ep_lesionnum_afterxpoly = sapply(upstream_grid, function(u){
      epcml_sitel_afterx =  mat_ep_lesionnum_detinitial[x,1:(Lmax+1)]%*% sapply(0:Lmax,function(b) dpoi
      return(epcml_sitel_afterx+(1-pv)*u)
    })

    return(ep_lesionnum_afterxpoly )

  }) %>% do.call(rbind,.)

  return( mat_epcml_lesionnum )
}
```

### Observed vs expected lesion density over multiple genes including invisible lesions

If we suppose we have $k$ genes then the expected observed lesion count up to position $x$, summed over the $k$ genes is

$$\mathbb{E}[L_n^{(k)}] = k(\mu x(1-p_v) + \mathbb{E}[L_n]).$$

In the absence of repair the expected number of mutations is

$$k\mu x.$$

Often we work with the density in an increment between two points $x_{i-1}$ and $x_i$. Here the expected observed number of lesions over $k$ genes in the increment is

$$\mathbb{E}[L_n^{(k)}(x_i)] - \mathbb{E}[L_n^{(k)}(x_{i-1})] = k(\mu x_i(1-p_v) + \mathbb{E}[L_n(x_i)]) - k(\mu x_{i-1}(1-p_v) + \mathbb{E}[L_n(x_{i-1})]) \tag{6}$$
$$= k(\mu(1-p_v)(x_i - x_{i-1}) + \mathbb{E}[L_n(x_i)]) - \mathbb{E}[L_n(x_{i-1})]). \tag{7}$$

The expected number of lesions in the absence of repair in the increment is

$$k\mu(x_i - x_{i-1}).$$

Therefore, the observed over expected number of lesions in the increment is

$$1 - p_v + \frac{\mathbb{E}[L_n(x_i)]) - \mathbb{E}[L_n(x_{i-1})]}{\mu(x_i - x_{i-1})}. \tag{8}$$

In R code this can be implemented as:

```
get_obsep_denslesions_MC = function(pd,
                                    pr,
                                    npoly,
                                    upstream_grid,
                                    Lmax,
                                    pv){

  mat_epcml_lesionnum = get_cmllesions_MC (pd,
                                           pr,
                                           npoly,
                                           upstream_grid,
                                           Lmax,
                                           pv)


  mat_obsdens_lesionnum =  lapply(1:nrow(mat_epcml_lesionnum), function(x){

    return(c(mat_epcml_lesionnum[x,1],diff(mat_epcml_lesionnum[x,])))

  }) %>% do.call(rbind,.)

  mat_obsepdens_lesionnum = mat_obsdens_lesionnum/(upstream_grid[2]-upstream_grid[1])

  return(   mat_obsepdens_lesionnum  )
}
```

We use Equation (8) to compare against the experimental data. The number of polymerases which pass through a gene $n$ is unknown; hence we introduced the expression multiplier $m$ such that $n = \text{round}(em)$, where $e$ is the average nascent transcript levels of the genes under consideration. By minimising the mean squared distance between Equation (8) and the data, the parameters $p_d$, $p_r$, $p_v$, $m$ may be inferred. The full code used for parameter inference is provided online, however after this point the inference is standard.
